# Kinetochore clustering is mediated by Mps1 phosphorylation of conserved MELT motifs in Stu1

**DOI:** 10.64898/2026.01.23.701336

**Authors:** Darren R. Mallett, Mengqiu Jiang, Gianna M. Minnuto, Sue Biggins

**Affiliations:** Molecular and Cellular Biology Graduate Program, University of Washington, 1705 NE Pacific Street, Seattle, WA 98195, USA; Howard Hughes Medical Institute, Division of Basic Sciences, Fred Hutchinson Cancer Center, 1100 Fairview Ave. N, Seattle, WA, 98109, USA

**Keywords:** kinetochore, Mps1 kinase, MELT motif, Stu1/CLASP, kinetochore clustering, spindle assembly checkpoint

## Abstract

Unattached kinetochores promote microtubule capture while preventing cell cycle progression during mitosis. The Mps1 kinase controls these events by mediating kinetochore assembly of the fibrous corona in animal cells and by triggering the spindle checkpoint. In budding yeast, which does not assemble a fibrous corona, the Stu1 and Slk19 spindle proteins promote microtubule capture by clustering unattached kinetochores, but the underlying mechanism is unclear. Here, we show that Mps1 controls this pathway by phosphorylating two conserved MELT motifs in Stu1 to recruit Slk19 and mediate kinetochore clustering. Structural analysis of the Stu1:Slk19 complex reveals long, string-like filaments and offers mechanistic insight into how kinetochores might cluster. Our findings suggest the regulation of kinetochore capture is a conserved Mps1 function across eukaryotes.

**SUMMARY:** Unattached kinetochores cluster in budding yeast to promote microtubule capture. Mallett et al. dissect the molecular mechanisms of this pathway, showing Mps1 kinase regulates an interaction between Stu1 and Slk19 to drive clustering.

## INTRODUCTION

During mitosis, cells undergo precise chromosome segregation to ensure that each daughter cell receives a complete copy of the genome. Chromosome segregation requires kinetochores, complex protein machines that assemble on centromeric chromatin, and microtubules, dynamic polymers that pull sister chromatids apart^1^. To ensure accurate segregation, every pair of sister kinetochores must biorient by attaching to microtubules from opposite poles. Once all sister kinetochores are bioriented, the cell triggers anaphase and pulls the sister chromatids apart^2^.

Kinetochores achieve biorientation through a complicated trajectory that initially involves making a lateral attachment to a microtubule prior to an end-on attachment^3–6^. Initial microtubule attachment is assisted by an oligomeric protein network recruited to the outer kinetochore termed the fibrous corona in metazoa whose assembly is initiated by Mps1 kinase activity^7–13^. Defects in kinetochore-microtubule attachments trigger the spindle checkpoint to prevent cell cycle progression until all chromosomes are bioriented, a conserved signaling cascade also controlled by Mps1^14–19^. Thus, Mps1 activity integrates the microtubule attachment machinery with the cell cycle machinery to promote attachments and prevent catastrophic errors.

In the budding yeast *Saccharomyces cerevisiae,* which lacks the corona pathway, microtubule capture is facilitated by the clustering of unattached kinetochores^20^. In this mechanism, a microtubule may capture a cluster of kinetochores in a single capturing event and transport them toward the spindle axis as a single unit. Consistent with this, the capture process and mitosis are prolonged in mutants defective in kinetochore clustering^20^. Stu1 and Slk19 are two proteins known to be involved in this pathway^20, 21^.

Stu1, also known as CLASP, is a microtubule rescue factor that is required for bipolar spindle formation^22–24^ and Slk19 is a multi-faceted protein with roles in spindle stability, centromere elasticity, and anaphase entry through a signaling network known as FEAR^25–28^. Similar to the corona and spindle checkpoint proteins, the Stu1 and Slk19 proteins robustly localize to unattached kinetochores, and Stu1 localization to unattached kinetochores requires Mps1 activity^21, 29, 30^. This suggests the role of Mps1 in promoting initial kinetochore-microtubule attachments is broadly conserved across eukaryotes.

In addition to kinetochore clustering, Stu1 and Slk19 support initial kinetochore capture in at least two other ways. Firstly, when unattached kinetochores are present, Stu1 and Slk19 relocate from the spindle axis to unattached kinetochores, an event that reorganizes spindle microtubules and promotes their outward growth into the nuclear space to “search” for unattached kinetochores^21^. Secondly, the presence of Stu1 and Slk19 at unattached kinetochores helps to stabilize the capturing microtubule to prevent kinetochore detachment, presumably through Stu1’s rescue function^21^.

It is unclear how Stu1 and Slk19 drive the clustering of unattached kinetochores and whether the spindle checkpoint is also involved in clustering. Here, we show that Mps1 controls unattached kinetochore clustering in a pathway independent of the checkpoint.

Our analyses reveal that Stu1 contains MELT motifs that are phosphorylated by Mps1 to directly recruit Slk19, which in turn promotes maximal recruitment of the complex *in vivo* and controls clustering. Structural analyses demonstrate that the Stu1:Slk19 complex forms >100 nm long filaments *in vitro*, offering a potential mechanism of how the complex may drive unattached kinetochore clustering in cells. Overall, our work underscores Mps1 as a critical regulator of unattached kinetochores where it promotes microtubule capture in budding yeast through the Stu1:Slk19 complex in addition to its established roles in spindle checkpoint signaling.

## RESULTS

### Stu1’s TOG1 domain contains a basic patch required for kinetochore binding

We set out to better understand the mechanism of unattached kinetochore clustering. We first tested whether Stu1 must localize to the kinetochore to mediate clustering. To do this, we sought to identify a specific Stu1 mutant that abolishes its kinetochore association. Stu1 is organized into 4 globular domains and 2 large, disordered regions (**Figure 1A**)^30^. Unlike traditional TOG domains, TOG1 does not bind tubulin dimers but is instead required for Stu1’s kinetochore recruitment^24, 30^. It was also reported that the disordered C-terminal loop (CL) region is required for Stu1’s localization to unattached kinetochores in live cell imaging assays^21, 30^, and it is also required for Stu1 to co-immunoprecipitate with Slk19^31^. To further dissect the functional differences between TOG1 and the CL region, we turned to biochemical methods. We immunoprecipitated FLAG-tagged Stu1 constructs lacking either TOG1 (*stu1^ETOG^*^1^) or the CL region (*stu1^ECL^*) from cells arrested with the microtubule destabilizing drug benomyl to generate unattached kinetochores and performed immunoblotting on Slk19 and kinetochore components (**Figure 1A**). We confirmed a previous report that full-length Stu1 copurifies Slk19^31^, and we found it also copurifies with the kinetochore protein Ndc80 and the spindle checkpoint component Bub1 (**Figure 1A**). Deletion of the TOG1 domain abolished the Stu1 interaction with Ndc80 and Bub1 while deletion of the CL region abolished its interaction with Slk19 but did not affect the interaction with Ndc80 and Bub1. Surprisingly, the TOG1 deletion also severely reduced the Slk19 interaction. Together, these biochemical results show that TOG1, rather than the CL region, is the kinetochore binding domain despite similar localization defects reported in fluorescence microscopy assays^21, 30^, and that both TOG1 and the CL are involved in Slk19 association.

**Figure 1:**
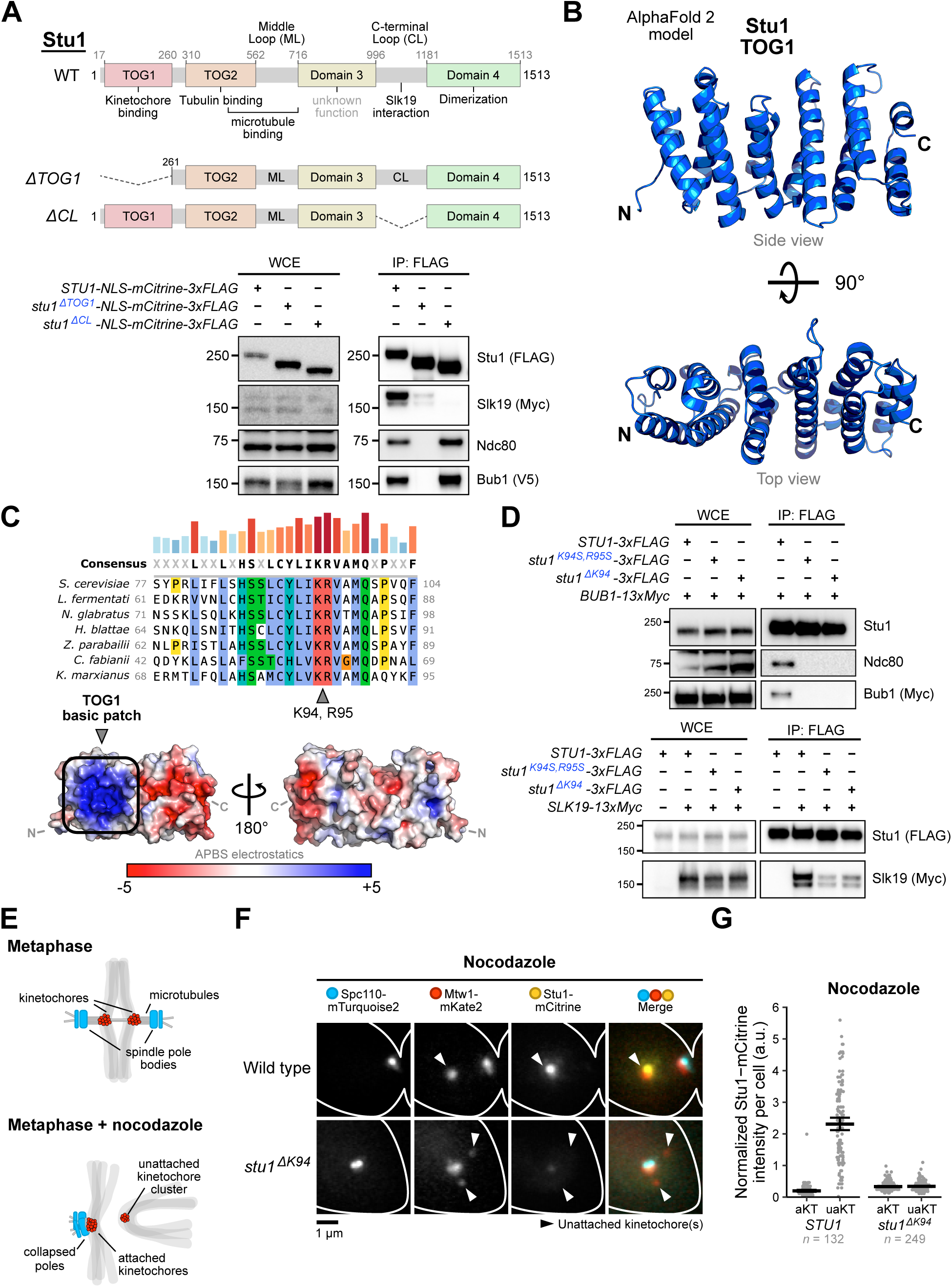
Analysis of Stu1’s TOG1 domain reveals a basic patch required for kinetochore binding. (A) Deletion of the Stu1 TOG1 domain abolishes its kinetochore interaction and decreases its Slk19 interaction, whereas deletion of the CL region abolishes its Slk19 interaction. Top: Stu1’s domain structure (as described in ^30^) with the indicated deletion constructs below the wild type (WT). Bottom: lysates from *STU1-NLS-mCitrine-3xFLAG* (SBY22069), *stu1^ETOG1^-NLS-mCitrine-3xFLAG* (SBY22050), and *stu1^ECL^-NLS-mCitrine-3xFLAG* (SBY22056) cells also containing *BUB3-3xV5* and *SLK1S-13xMyc* were immunoprecipitated using α-FLAG beads and analyzed by immunoblotting with the indicated antibodies. Note that Slk19-13xMyc has two bands: the full-length protein and a smaller cleavage product^76^. **(B)** AlphaFold 2 model of Stu1’s TOG1 domain (residues 17-260, ribbon diagram) shows a paddle-like structure consisting of HEAT repeats as described for other TOG domains^33, 34^. **(C)** *S. cerevisiae* Stu1 residues K94 and R95 are conserved and contribute to a basic patch on the surface of TOG1. Top: multiple sequence alignment of a short region belonging to the TOG1 domain with sequences from several yeasts of the *Saccharomycetes* class (budding yeasts). The bars above the consensus sequence are a visual aid of conservation (blue, shorter: less conserved; red, taller: more conserved). Bottom: surface electrostatic potential analysis of the TOG1 AlphaFold 2 model reveals a densely packed basic patch where K94 and R95 are situated. **(D)** Mutations in TOG1’s basic patch disrupt kinetochore binding and reduce the interaction with Slk19. Lysates from benomyl-arrested *STU1-3xFLAG BUB1-13xMyc* (SBY21197), *stu1^KS4S, RS5S^-3xFLAG BUB1-13xMyc* (SBY21810), and *stu1^EKS4^-3xFLAG BUB1-13xMyc* (SBY21811) cells (top) and *STU1-*3xFLAG (SBY21199), STU1*-3xFLAG SLK1S-13xMyc* (SBY21979), *stu1^KS4S, RS5S^-3xFLAG SLK1S-13xMyc* (SBY21943), and *stu1^EKS4^-3xFLAG SLK1S-13xMyc* (SBY21935) cells (bottom) were immunoprecipitated using α-FLAG beads and analyzed by immunoblotting with the indicated antibodies. (E-G) Mutating TOG1’s basic patch disrupts Stu1-mCitrine localization to unattached kinetochores in live cell imaging assays. **(E)** Schematic showing kinetochore and spindle pole behavior in *S. cerevisiae* cells during metaphase with and without nocodazole treatment. Top: kinetochores form a bilobed configuration by attaching to microtubules from opposite poles. Bottom: when treated with nocodazole, most kinetochores remain attached to spindle poles while a subset of kinetochores detach from spindle poles and cluster together^20, 36^. **(F)** Stu1-mCitrine (SBY21184 cells) and Stu1^ΔK94^-mCitrine (SBY24733 cells) were visualized on unattached kinetochores in cells also expressing *SPC110-mTurquoise2* and *MTW1-mKate2* using live cell imaging. Cells were released from G1 into 15 µg/mL nocodazole for 1.5 hours to create unattached kinetochores before imaging. A single representative cell is shown for each genotype (rows), and each fluorescent channel is indicated above. **(G)** Quantification of mCitrine fluorescence of the experiment shown in (F). mCitrine signals were measured at both attached and unattached kinetochores (aKTs and uaKTs, respectively) and reported as an Mtw1-normalized ratio (mCitrine fluorescence/mKate2 fluorescence). Each dot represents the fluorescence value within a single cell. Thick horizontal bars represent the mean value, and error bars represent the 95% confidence interval as determined by bootstrapping, the values of which are shown in **Table S4**. (**A and D**) WCE, whole-cell extract.

We next set out to identify a specific TOG1 mutant that abolishes Stu1 kinetochore binding while keeping the domain largely intact. The AlphaFold 2^32^ model of TOG1 shows a classic TOG-like paddle structure with 6 HEAT repeats^33, 34^ (**Figure 1B**). A multiple sequence alignment with other *Saccharomycetes* yeast species shows clear stretches of residues with high sequence conservation throughout TOG1 (**Figure 1C, Figure S1A**). We hypothesized that TOG1, like spindle checkpoint proteins, possibly recognizes a phosphorylated interface on the kinetochore^18, 35^. Analysis of TOG1’s surface charge distribution revealed a conserved, densely packed ‘basic patch’ on one side of the paddle structure towards the N-terminus that is suitable for electrostatic binding with a phosphorylated receptor (**Figure 1C, Figure S1B**). To disrupt the basic patch, we targeted two conserved residues, K94 and R95, using CRISPR-Cas9 mutagenesis (**Figure 1C**). The mutagenesis resulted in two separate mutants: (1) K94S, R95S and (2) ΔK94, in which K94 was deleted while R95 remained intact. Immunoprecipitation of both mutants mirrored that of Stu1^ΔTOG1^ in which the interactions with the kinetochore proteins Ndc80 and Bub1 were abolished while the interaction with Slk19 was severely reduced (**Figure 1D**). Both mutants displayed increased fitness in the presence of the microtubule-destabilizing drug benomyl for unknown reasons compared to wild type cells but otherwise displayed normal growth under standard conditions (**Figure S1C**). Although the Stu1^ΔK94^ mutant is likely to be more structurally disruptive, silver stain analysis of the mutant proteins indicated that the double serine mutant (Stu1^K94S, R95S^) may be less stable because there were more degradation products compared to Stu1^ΔK94^ (see degradation bands, **Figure S1D**). We therefore moved forward with studying the Stu1^ΔK94^ mutant.

We tested whether the *stu1^EKS4^* mutation disrupts Stu1’s localization to unattached kinetochores using live cell imaging. Budding yeast kinetochores are clustered throughout most of the cell cycle regardless of their attachment status^20, 36–38^. During a nocodazole arrest (which depolymerizes microtubules), the spindle pole bodies collapse, and kinetochores either remain co-localized with the collapsed poles or localize distally^20, 21, 36^ (**Figure 1E**). Prior work established that the kinetochores that cluster with the collapsed poles remain attached to short microtubules^20, 36^. Therefore, spindle checkpoint proteins only localize to the unattached kinetochore clusters that are distal from the poles^36^. Thus, the attachment status of a particular kinetochore cluster can be determined using either a spindle checkpoint marker or a spindle pole body marker. We used the latter to ensure the nocodazole treatment was effective and the poles collapsed. We C-terminally tagged the spindle pole body protein Spc110, the kinetochore protein Mtw1, and wild type Stu1 or Stu1^ΔK94^ with mTurquoise2, mKate2, and mCitrine, respectively, and performed live cell imaging after release from a G1 arrest with or without nocodazole. In unperturbed metaphase cells, Stu1-mCitrine localized in between the spindle pole bodies along the spindle axis as previously described^29, 30^ (**Figure S1E**). The localization of Stu1^ΔK94^-mCitrine was similar to wild type in an unperturbed metaphase, confirming the point mutation does not affect Stu1’s spindle localization (**Figure S1E**). In nocodazole-treated cells, Stu1-mCitrine robustly localized to unattached kinetochores as expected^21, 29^ (**Figure 1F and 1G**). In contrast, Stu1^ΔK94^-mCitrine no longer localized to unattached kinetochores, confirming our biochemical data (**Figure 1F and 1G**). Taken together, we identified a specific mutant in Stu1’s TOG1 basic patch that abolishes Stu1’s interaction with the kinetochore.

### *stu1* TOG1 mutants are defective in unattached kinetochore clustering and Slk1G recruitment

We next tested whether Stu1 binding to unattached kinetochores is required for clustering and how that compares to the clustering defects of *slk1S*Δ cells. Wild type, *slk1S*Δ, and *stu1^EKS4^* cells containing the bright kinetochore marker Mtw1-3xmYPet were synchronized in G1 and released into media containing nocodazole to generate unattached kinetochores (**Figure 2A**). As previously observed, wild type cells often had two to three kinetochore foci, indicating robust clustering of the 16 sister kinetochores, while *slk1S*Δ cells exhibited kinetochore declustering (2.61 kinetochore foci per cell for wild type vs. 4.83 foci per cell for *slk1S*Δ, *p* < 0.001, **Figure 2B**)^20^. *Stu1^EKS4^* cells also displayed kinetochore declustering compared to wild type controls, showing Stu1’s kinetochore binding is required for clustering (2.68 kinetochore foci per cell for *STU1-3xFLAG* vs. 3.98 foci per cell for *stu1^EKS4^-3xFLAG, p* < 0.001, **Figure 2B**). Since *SLK1S* is required for clustering, we subsequently tested whether Slk19-mCitrine could localize to unattached kinetochores in *stu1^EKS4^* cells (**Figure 2C**). *Stu1^EKS4^* cells showed a 76% reduction in Slk19-mCitrine fluorescence at unattached kinetochores compared to wild type controls (**Figure 2D**). Taken together, these results show that the recruitment of Stu1 to unattached kinetochores through its TOG1 domain promotes Slk19 recruitment and is required for robust kinetochore clustering.

**Figure 2:**
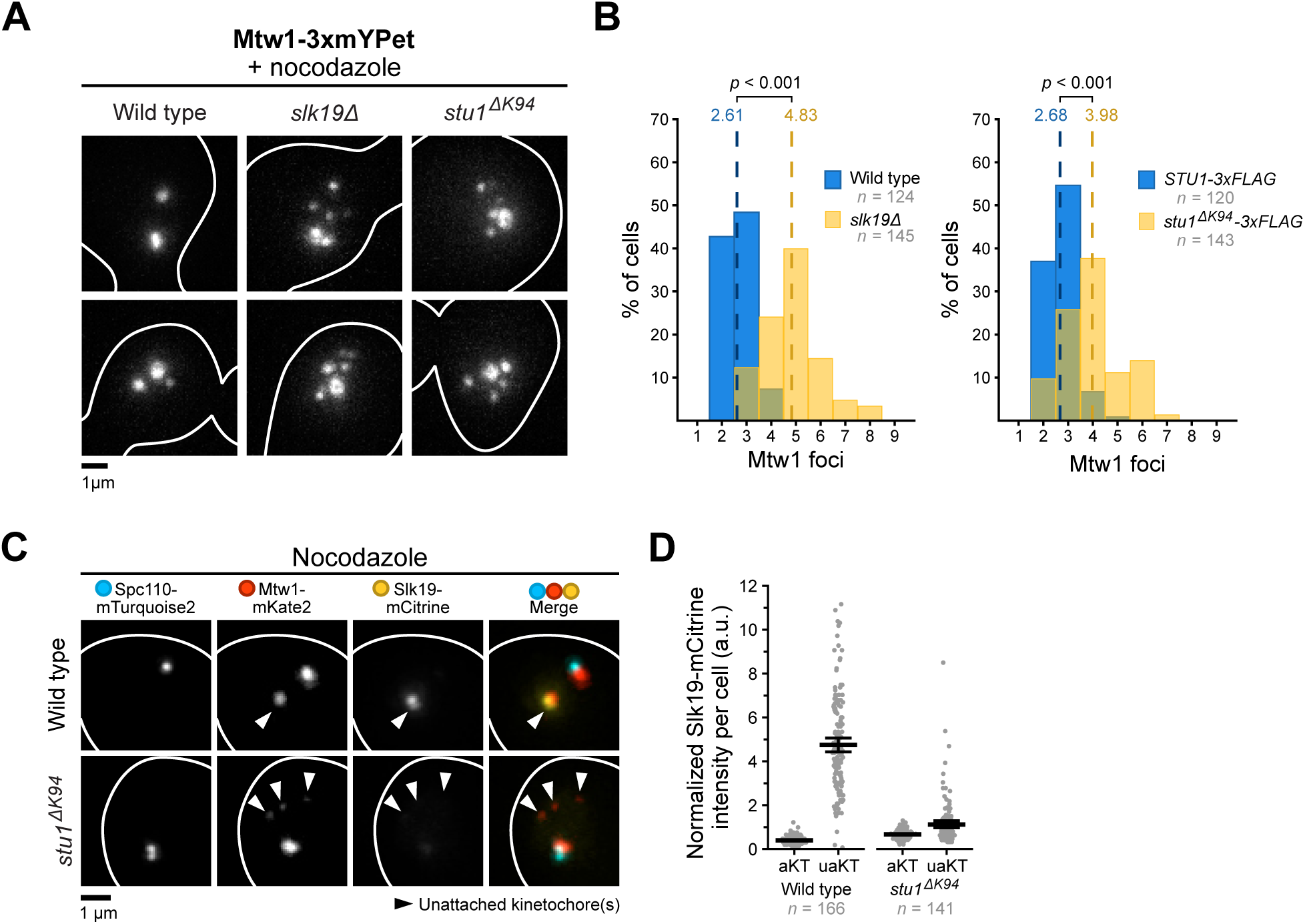
Stu1 TOG1 mutants display kinetochore clustering defects and altered Slk1G localization. (A) *Stu1^EKS4^* cells have clustering defects. The degree of unattached kinetochore clustering was assessed using *MTW1-3xmYPet* in wild type (SBY23480), *slk1S*Δ (SBY23372), and *stu1^EKS4^*(SBY23484) cells. Cells were released from G1 into 15 µg/mL nocodazole for 1.5 hours to create unattached kinetochores and live imaged. Each genotype shows two representative cells. **(B)** Quantification of Mtw1-3xmYPet foci counts per cell from the experiment in (A). Dashed lines represent the mean foci count for each strain. *p-*values were generated using a Mann-Whitney U test. **(C)** *Stu1^EKS4^* disrupts Slk19-mCitrine localization to unattached kinetochores. Slk19-mCitrine was visualized on unattached kinetochores in *STU1* (SBY22043) and *stu1^EKS4^* (SBY21963) mutants containing fluorescent Spc110-mTurquoise2 and Mtw1-mKate2. Cells were released from G1 into 15 µg/mL nocodazole for 1.5 hours to create unattached kinetochores and live imaged. A representative cell is shown for each genotype, and each fluorescent channel is indicated above. **(D)** Quantification of mCitrine fluorescence intensity at attached and unattached kinetochores (aKT and uaKT, respectively) from the experiment shown in (C). mCitrine fluorescence intensities were measured and reported as in Figure 1G. Thick horizontal bars represent the mean value, and error bars represent the 95% confidence interval as determined by bootstrapping, the values of which are shown in **Table S4**.

### The Stu1-Slk1G interaction is promoted at unattached kinetochores

Because the Stu1 mutant that does not localize to kinetochores also alters Slk19 localization, we hypothesized that their interaction is promoted at the kinetochore. This would explain the reduction in the Stu1-Slk19 interaction when TOG1 is deleted or mutated (**Figure 1A and 1D**). To test this, we asked whether the Stu1-Slk19 interaction could be rescued if we forced the Stu1 TOG1 mutant protein to localize to unattached kinetochores. The spindle checkpoint protein Bub3 robustly binds unattached kinetochores through Mps1 phosphorylation of Spc105’s MELT motifs^16, 35^. We therefore fused Bub3 to the N-terminus of the Stu1 TOG1 mutant (Bub3-Stu1^ΔK94^-3xFLAG) and analyzed the levels of co-purifying Slk19 when cells were treated with the microtubule destabilizing drug benomyl. Supporting our hypothesis, the Bub3-Stu1^ΔK94^-3xFLAG fusion protein restored co-purifying Slk19 levels similarly to wild type controls (**Figure 3A**). The kinetochore association of the Stu1 TOG1 mutant protein was also restored as shown by its interaction with the Ndc80 and Spc105 kinetochore proteins (**Figure 3A**).

**Figure 3:**
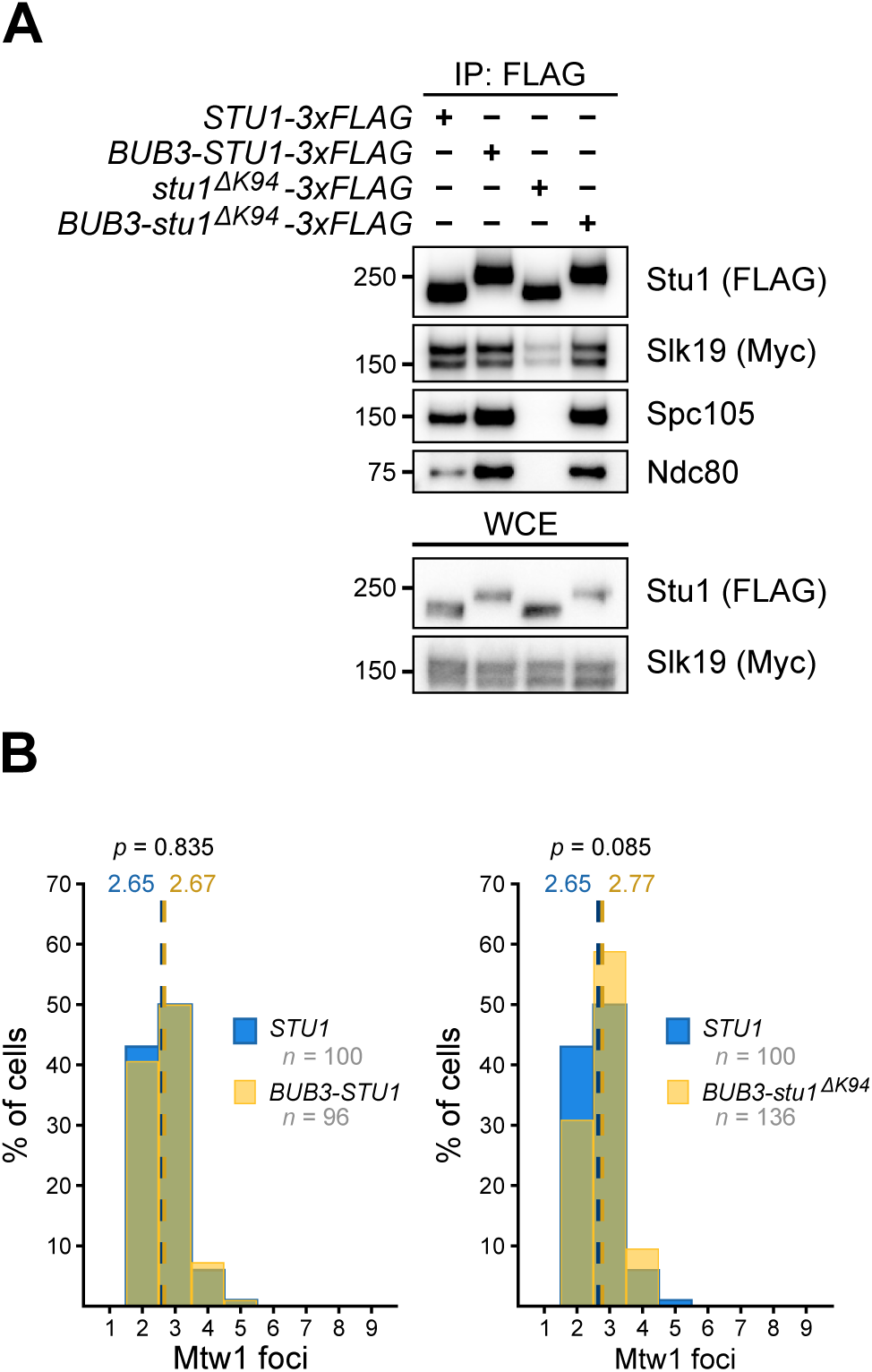
The Stu1-Slk1G interaction is promoted at kinetochores. (A) Fusing Bub3 to the N-terminus of Stu1^ΔK94^ rescues the levels of co-purifying Slk19. *STU1-3xFLAG* (SBY21979), *BUB3-STU1-3xFLAG* (SBY23897), *stu1^EKS4^-3xFLAG* (SBY21935), and *BUB3-stu1^EKS4^-3xFLAG* (SBY23901) cells containing *SLK1S-13xMyc* were arrested in benomyl and lysates were immunoprecipitated using α-FLAG beads and analyzed by immunoblotting using the indicated antibodies. WCE, whole-cell extract. **(B)** *BUB3-stu1^EKS4^* rescues the clustering defects of *stu1^EKS4^* cells. *MTW1-3xmYPet* cells containing *STU1* (SBY23480), *BUB3-STU1* (SBY24036), or *BUB3-stu1^EKS4^* (SBY24038) were released from G1 into 15 µg/mL nocodazole for 1.5 hours to create unattached kinetochores and live imaged. Graphs display the quantification of Mtw1-3xmYPet foci counts per cell. *p-*values were generated using a Mann-Whitney U test. The wild type data in both graphs (*STU1*, blue) are the same for easier comparison.

We next tested whether the fusion protein rescues the kinetochore clustering defects observed in *stu1^EKS4^* cells. As a control, we confirmed that the fusion of Bub3 to wild type Stu1 does not affect clustering in cells treated with nocodazole (**Figure 3B**, *STU1* vs. *BUB3-STU1*, *p* = 0.835). In alignment with our hypothesis, cells expressing *BUB3-stu1^EKS4^* exhibited normal kinetochore clustering (**Figure 3B**, *STU1* vs. *BUB3-stu1^EKS4^, p* = 0.085), unlike *stu1^EKS4^* cells as shown above. Taken together, these data strongly suggest that Stu1 must bind to unattached kinetochores through its TOG1 domain to efficiently interact with Slk19 to promote clustering.

### The spindle checkpoint and unattached kinetochore clustering are separate pathways controlled by Mps1

The kinetochore localization of Stu1 and Slk19 resembles that of spindle checkpoint complexes that require Mps1 to specifically mediate their interaction at unattached kinetochores^16, 18^. In addition, Stu1 fails to localize to unattached kinetochores in cells inhibited for Mps1 activity^21^. However, it has not been tested whether Mps1 or other spindle checkpoint proteins are required for unattached kinetochore clustering. Furthermore, it is unclear whether clustering affects spindle checkpoint function. We therefore set out to understand the relationship between Mps1, unattached kinetochore clustering, and the spindle checkpoint (**Figure 4A**).

**Figure 4:**
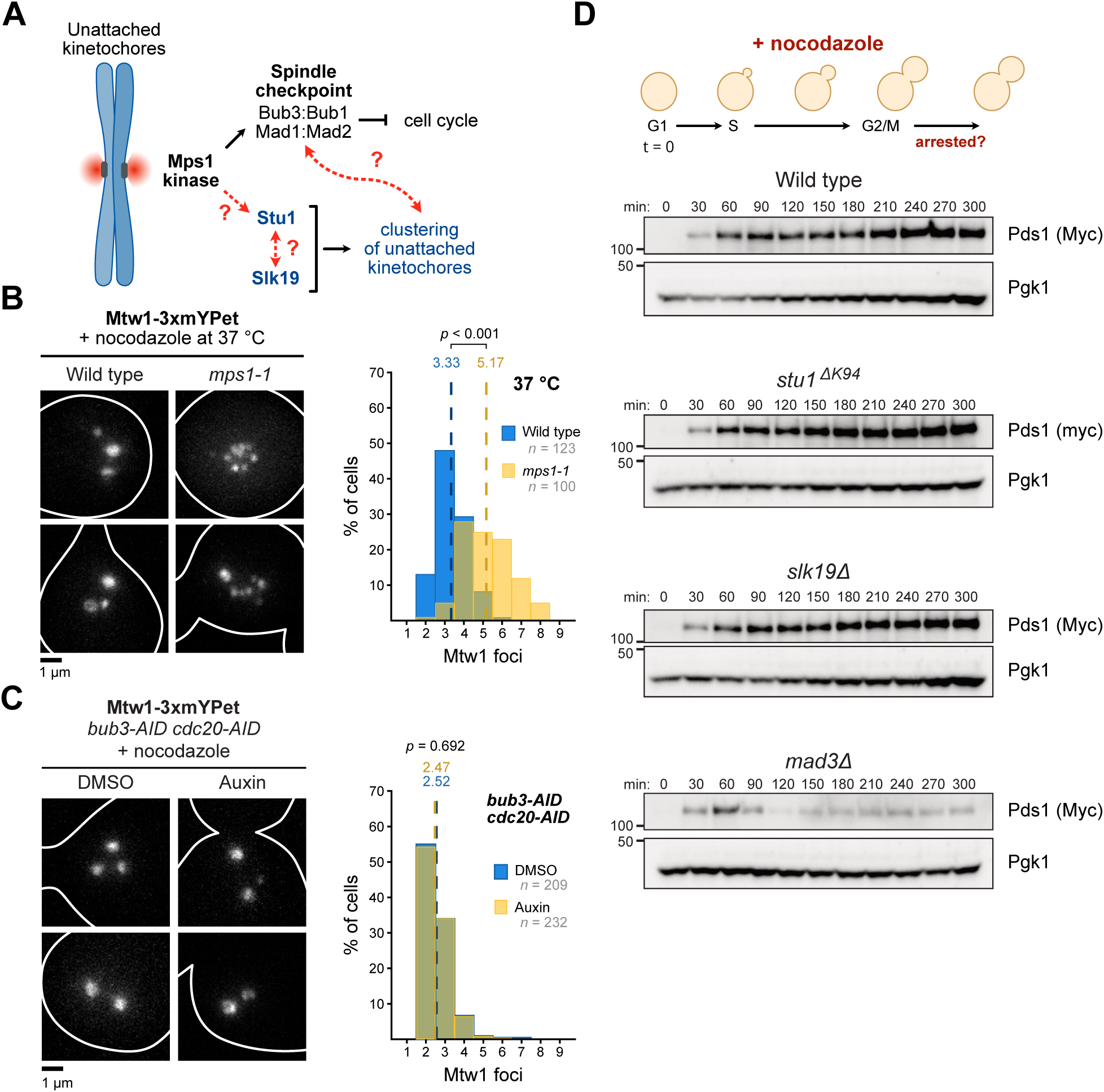
Mps1 separately controls the spindle checkpoint and unattached kinetochore clustering. **(A)** Schematic displaying unknown relationships between Mps1, Stu1, Slk19, and the spindle checkpoint. **(B)** Mps1 is required for kinetochore clustering. Wild type (SBY23370) and *mps1-1* (SBY23478) cells containing fluorescent Mtw1-3xmYPet were arrested in G1 and released into 15 µg/mL nocodazole at the restrictive temperature of 37 °C. Left: two representative cells are shown for each genotype. Right: quantification of Mtw1-3xmYPet foci counts per cell. Dashed lines represent the mean foci count for each strain. *p-*values were generated using a Mann-Whitney U test. **(C)** The spindle checkpoint is not required for unattached kinetochore clustering. Unattached kinetochores were generated in *MTW1-3xmYPet bub3-AID cdc20-AID* cells (SBY23547) similarly to (B), except 2 mM auxin (or equal volume of DMSO as a control) was added to degrade Bub3 and Cdc20. Left: two representative cells are shown for each condition. Right: quantification of kinetochore foci counts is shown following the same methods as (B). **(D)** Disrupting unattached kinetochore clustering does not affect spindle checkpoint function. Wild type (SBY22005), *stu1^EKS4^* (SBY21961), *slk1S*𝛥 (SBY8968), and *mad3*𝛥 (SBY22334) cells expressing *PDS1-18xMyc* were arrested in G1, released into media containing 15 µg/mL nocodazole, and the levels of Pds1-18xMyc in whole-cell extracts were monitored at the indicated time points by immunoblotting using the indicated antibodies. α-Pgk1 was used as a loading control.

We asked whether the *mps1-1* temperature sensitive mutant^39^ displays unattached kinetochore clustering defects, as would be expected if Mps1 is required for Stu1 localization^21^. We released wild type and *mps1-1* cells from a G1 arrest into nocodazole at the nonpermissive temperature of 37 °C to generate unattached kinetochores and inhibit Mps1 activity (**Figure 4B**). Compared to wild type, *mps1-1* cells showed significantly increased numbers of kinetochore foci, with an average of ∼5 kinetochore foci per cell and a wider distribution (wild type vs. *mps1-1* at 37°C, *p* < 0.001, **Figure 4B**). These data confirm that *MPS1* is required for unattached kinetochore clustering.

We sought to determine whether the spindle checkpoint and kinetochore clustering pathways are interconnected or strictly separate. We tested whether abolishing the spindle checkpoint affects kinetochore clustering using a Bub3 auxin-inducible degron (*bub3-AID*)^35, 40, 41^. To retain a metaphase arrest in *bub3-AID* cells in the presence of nocodazole, we also included a *cdc20-AID* allele that blocks anaphase progression^42^. There was no difference in the number of kinetochore foci in cells depleted for Bub3 and Cdc20 compared to mock controls, indicating spindle checkpoint activation is not involved in kinetochore clustering (2.52 foci per cell in DMSO vs. 2.47 foci per cell in auxin, *p* = 0.692, **Figure 4C**). These data agree with previous findings showing Bub3 and the downstream spindle checkpoint proteins are not required for Stu1 localization to unattached kinetochores^21, 29^.

We next asked whether kinetochore clustering is required for spindle checkpoint activation. We examined checkpoint activation in *slk1S*Δ and *stu1^EKS4^* cells compared to controls by monitoring the levels of the anaphase inhibitor Pds1 after releasing cells from a G1 arrest into nocodazole^43^. Similar to wild type, *slk1S*Δ and *stu1^EKS4^* cells activated the spindle checkpoint and stabilized Pds1. As a control, spindle checkpoint-deficient *mad3*𝛥 cells did not arrest, indicated by Pds1 degradation between 60 and 90 minutes after release. These data suggest the clustering of unattached kinetochores is not required for spindle checkpoint activation (**Figure 4D**). Overall, our results are consistent with a model in which there is a bifurcation in Mps1 function at unattached kinetochores to regulate spindle checkpoint activation and kinetochore clustering independently.

### The Stu1-Slk1G interaction requires Mps1 and phosphorylation of Stu1’s MELT motifs

We next set out to determine how the Stu1-Slk19 interaction is promoted at unattached kinetochores. Since their behavior at unattached kinetochores resembles spindle checkpoint complexes^16, 18, 21^, we asked whether their co-purification depends on Mps1. However, Stu1 and Slk19 are not known to be Mps1 substrates, so we also tested the polo kinase Cdc5 because it phosphorylates Slk19 and other spindle proteins^44^. We immunoprecipitated Stu1-3xFLAG from wild type, *mps1-1,* and *cdc5-1* cell lysates and found that Slk19 co-purified with Stu1 from wild type and *cdc5-1* cells but not from *mps1-1* cells at the restrictive temperature, indicating that Mps1 activity promotes the Stu1-Slk19 interaction (**Figure 5A**). These data suggested that phosphorylation is required for the Stu1-Slk19 interaction, so we tested this by immunoprecipitating Stu1-3xFLAG and treating the bead-bound proteins with 𝜆-phosphatase in the presence or absence of phosphatase inhibitors. Phosphatase treatment caused the co-purifying Slk19 to dissociate from the beads specifically in the absence of inhibitors, indicating a phospho-dependent interaction (**Figure 5B**).

**Figure 5:**
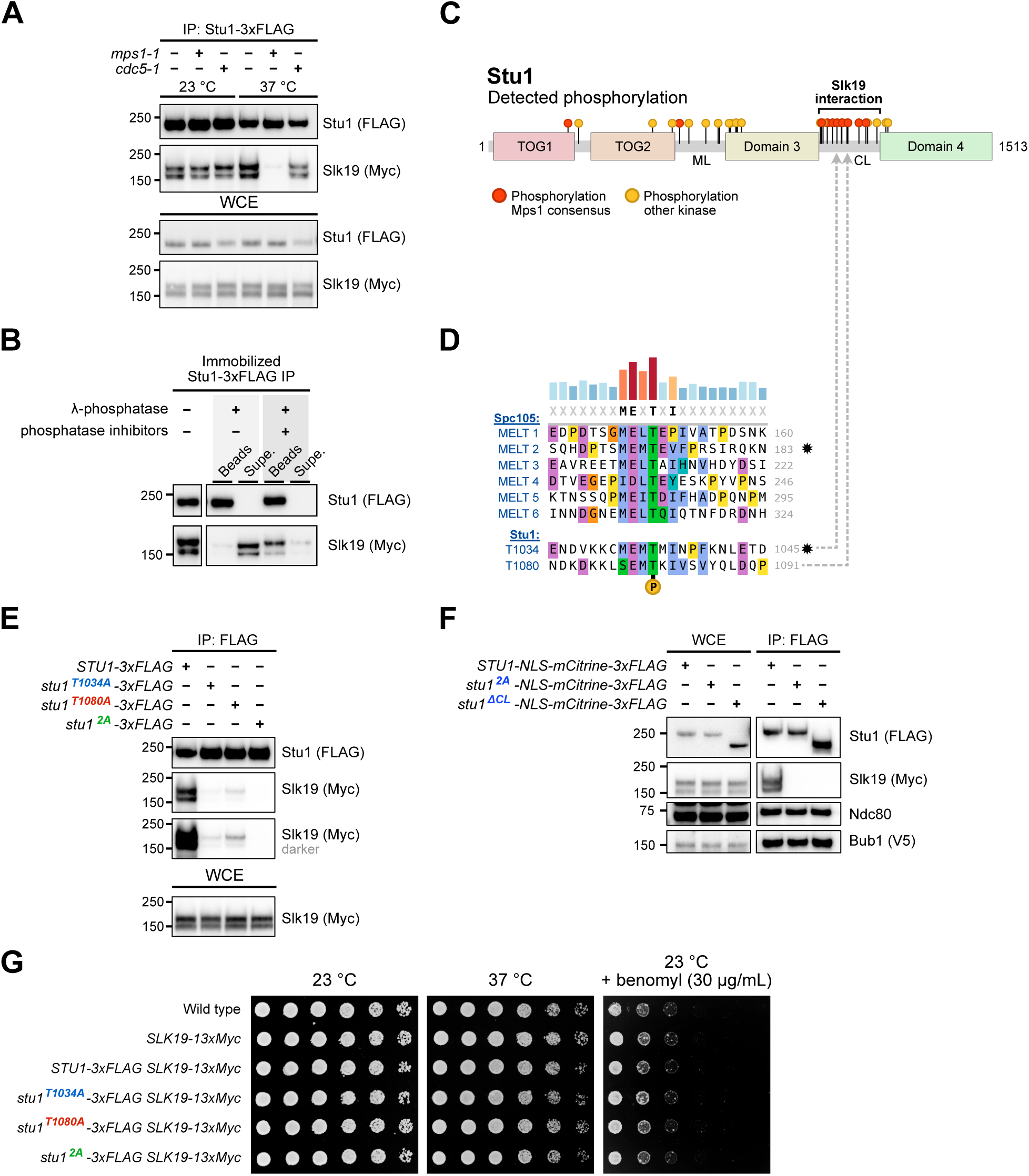
The Stu1-Slk1G interaction requires Mps1 and phosphorylation of Stu1’s MELT motifs. **(A)** Mps1 activity is required for the Stu1-Slk19 interaction. Stu1-3xFLAG was immunoprecipitated using α-FLAG beads from *STU1-3xFLAG SLK1S-13xMyc* (SBY21979), *STU1-3xFLAG SLK1S-13xMyc mps1-1* (SBY21981), and *STU1-3xFLAG SLK1S-13xMyc cdc5-1* (SBY22330) lysates grown at the indicated temperatures and analyzed by immunoblotting using the indicated antibodies. **(B)** Stu1 associates with Slk19 in a phospho-dependent manner. Stu1-3xFLAG was immunoprecipitated from *STU1-3xFLAG SLK1S-13xMyc* lysate (SBY21979) using α-FLAG beads and treated with λ-phosphatase (with or without phosphatase inhibitors). Bound and unbound (supernatant) fractions were analyzed by immunoblotting with the indicated antibodies. **(C)** Immunoprecipitates from *STU1-3xFLAG cdc20-AID leu2::pGPD1-OsTIR1* cells (SBY20662) were analyzed by mass spectrometry from 3 conditions: asynchronous (DMSO), benomyl-arrested (30 µg/mL benomyl; metaphase arrest, spindle checkpoint active), and *cdc20-AID* arrested (500 µM auxin IAA; metaphase arrested, spindle checkpoint inactive). The schematic displays the cumulative detected phosphorylation sites across all conditions (for individual conditions and coverage maps, see **Figure S3**). Sites in red follow the Mps1 consensus sequence, and the region required for the Slk19 interaction^31^ is highlighted. **(D)** Multiple sequence alignment of *S. cerevisiae* Spc105 MELT motifs with Stu1 MELT motifs. The MELT surrounding Stu1 T1034 is closely related to MELT 2 of Spc105 (stars). Dashed lines point to their position within the CL region shown in (C). **(E)** Preventing Stu1 MELT phosphorylation disrupts the Stu1-Slk19 interaction. Lysates from benomyl-arrested *STU1-3xFLAG* (SBY21979), *stu1^T1034A^-3xFLAG* (SBY23770), *stu1^T1080A^-3xFLAG* (SBY22476), and *stu1^2A^-3xFLAG* (SBY23751) cells containing *SLK1S-13xMyc* were immunoprecipitated using α-FLAG beads and analyzed by immunoblotting with the indicated antibodies. **(F)** The immunoprecipitation behavior of *stu1^2A^* and *stu1*^Δ^*^CL^* are identical. Lysates from benomyl-arrested *STU1-NLS-mCitrine-3xFLAG* (wild type, SBY22069), *stu1^2A^-NLS-mCitrine-3xFLAG* (SBY23112), and *stu1*^Δ^*^CL^-NLS-mCitrine-3xFLAG* (SBY22056) cells containing *SLK1S-13xMyc* and *BUB1-3xV5* were immunoprecipitated and analyzed as in (E). **(G)** Stu1 MELT mutants do not exhibit growth defects. Five-fold serial dilutions of wild type (SBY3), *SLK1S-13xMyc* (SBY8956), and *SLK1S-13xMyc* cells containing either the *STU1-3xFLAG* (SBY21979), *stu1^T1034A^-3xFLAG* (SBY23770), *stu1^T1080A^-3xFLAG* (SBY22476), or *stu1^2A^-3xFLAG* (SBY23751) allele were incubated at the indicated conditions for 24-72 hours and imaged after sufficient growth. (**A, E, and F**) WCE, whole-cell extract.

We sought to identify the specific phosphorylated residues required for the interaction. The region of Slk19 that interacts with Stu1 is unknown, so we focused on the Stu1 CL region and looked for phosphorylation sites in this region. We performed mass spectrometry on Stu1 purified from benomyl-arrested, metaphase-arrested, and asynchronous cultures. We detected phosphorylation sites along the length of Stu1, most of which concentrated within the middle loop (ML) and CL regions (**Figure 5C, Figure S2**), although these regions also received the most mass spectrometry coverage (**Figure S3**). To narrow down key sites, we identified sites that followed the Mps1 consensus sequence (**Figure 5C**)^45, 46^. Using a Stu1 sequence alignment with other budding yeasts, we found two heavily conserved phosphorylation sites in all three treatment conditions within Stu1’s CL region predicted to be Mps1 sites. Strikingly, these sites appear to be closely related to Spc105’s MELT motifs, Mps1 sites required for the initiation of spindle checkpoint signaling that directly recruit Bub3:Bub1^16, 35^ (**Figure 5D, Figure S2**). The first Stu1 MELT (1031-MEMT-1034) and its surrounding sequence is nearly identical to Spc105 MELT2, whereas the second Stu1 MELT (1077-SEMT-1080) appears to have diverged from all Spc105 MELTs in that it contains a serine instead of methionine or isoleucine at the start of the motif (**Figure 5D**). Interestingly, most of the other phosphorylation sites within Stu1 matching the Mps1 consensus clustered within the CL region, suggesting Mps1 may heavily target this region (**Figure 5C**). However, these additional sites in the CL are less conserved than the MELTs, so we focused our analysis on the MELTs (**Figure S2**).

To determine whether the MELTs are required for the Stu1-Slk19 interaction, we introduced phosphodeficient alanine substitutions at each site (*stu1^T1034A^* or *stu1^T1080A^*) or both (*stu1^T1034A, T1080A^* double mutant, hereafter referred to as *stu1^2A^*) and immunoprecipitated 3xFLAG-tagged mutant proteins. Preventing phosphorylation of either Stu1 residue T1034 or T1080 severely reduced the interaction with Slk19 while the Stu1^2A^ double mutant completely abolished it, implicating the MELTs in Slk19 binding (**Figure 5E**). To determine if the *stu1^2A^* mutant behaves the same as deleting the entire CL region (*stu1*^Δ^*^CL^*), we immunoprecipitated them and both mutants abolished the interaction with Slk19 while maintaining their interactions with the kinetochore protein Ndc80 and checkpoint protein Bub1 (**Figure 5F**). The phosphodeficient MELT mutants did not affect cell viability or growth compared to wild type cells, consistent with Stu1’s CL region and *SLK19* not being essential for viability^25, 30^ (**Figure 5G**). Unlike the *stu1* TOG1 mutants, the *stu1* MELT mutants did not show increased fitness when grown on benomyl, suggesting the increased fitness is related to a TOG1-specific function and not Stu1’s interaction with Slk19 (compare **Figure S1C** and **Figure 5G**). Taken together, these results show the Stu1-Slk19 interaction depends on phosphorylation of Stu1’s MELT motifs, and that these sites are likely regulated by Mps1.

### Stu1 MELT phosphorylation is required for Slk1G and Stu1 localization and affects kinetochore clustering

We next tested the function Stu1 MELT phosphorylation *in vivo*. First, we asked whether phosphorylation of the Stu1 MELT motifs recruits Slk19 to unattached kinetochores (**Figure 6A**). Compared to wild type, *stu1^T1034A^* and *stu1^T1080A^* cells showed a 55% and a 20% reduction in Slk19-mCitrine fluorescence at unattached kinetochores, respectively (uaKTs, **Figure 6B**). *Stu1^2A^* cells harbored a more drastic phenotype, showing an 83% reduction in Slk19-mCitrine levels at unattached kinetochores compared to wild type (**Figure 6B**). We also measured slight increases in Slk19-mCitrine fluorescence co-localizing with attached kinetochores in the mutants (aKTs, **Figure 6B**), although the fluorescence intensities were nowhere near the intensities of Slk19-mCitrine at unattached kinetochores in wild type cells.

**Figure 6:**
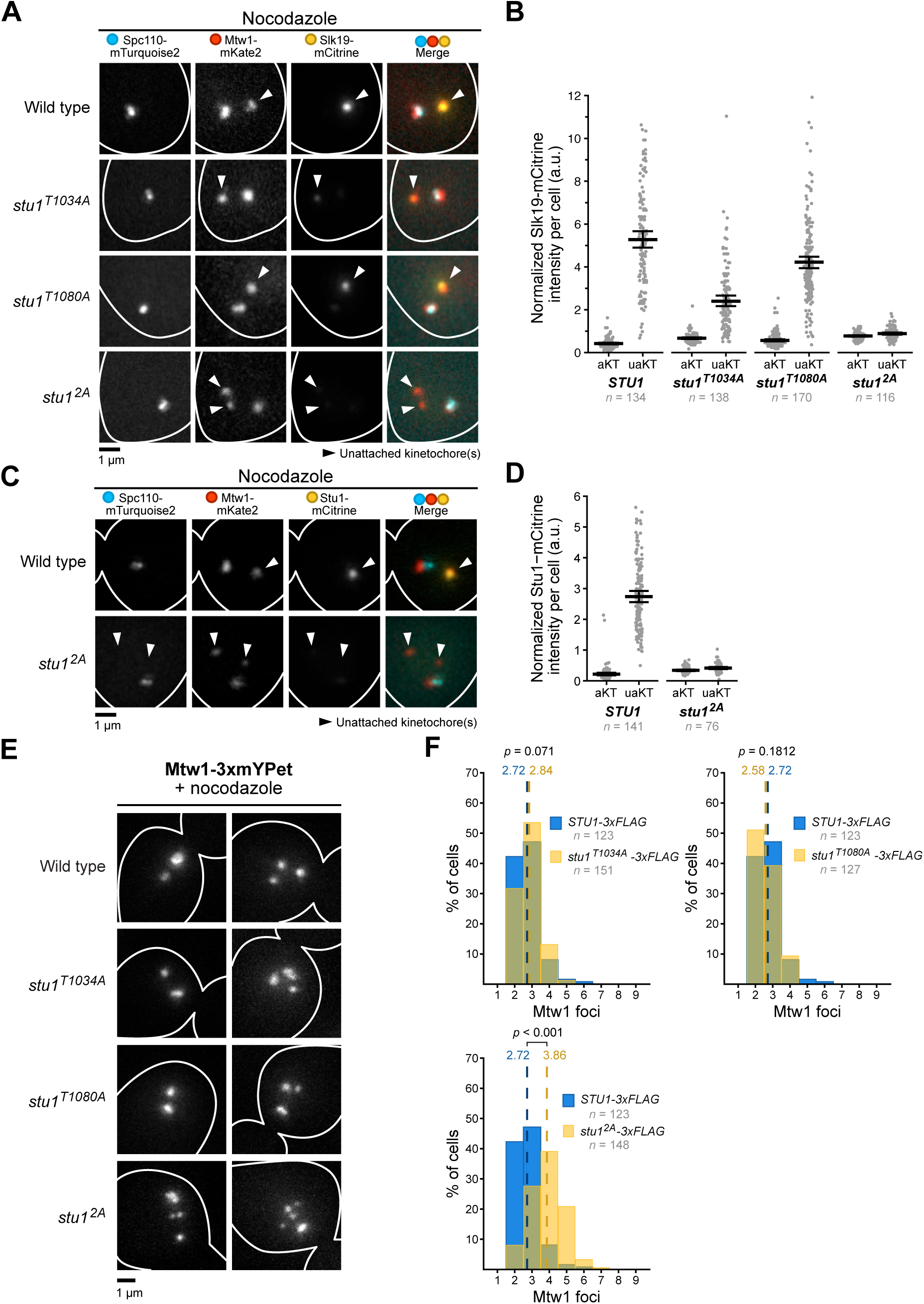
***Stu1^2A^* disrupts Slk1G and Stu1 localization and has kinetochore clustering defects. (A)** *Stu1* MELT mutants alter Slk19-mCitrine levels at unattached kinetochores. *STU1-3xFLAG* (wild type, SBY22043), *stu1^T1034A^-3xFLAG* (SBY24326), *stu1^T1080A^-3xFLAG* (SBY24331), and *stu1^2A^-3xFLAG* (SBY24246) cells containing fluorescent Spc110-mTurquoise2, Mtw1-mKate2, and Slk19-mCitrine were released from G1 into 15 µg/mL nocodazole for 1.5 hours to create unattached kinetochores and live imaged. A representative cell is shown for each genotype, and each fluorescent channel is indicated above. **(B)** Quantification of Slk19-mCitrine fluorescence from the experiment shown in (A). Slk19-mCitrine fluorescence was measured at both attached and unattached kinetochores (aKTs and uaKTs, respectively) and reported as an Mtw1-normalized ratio (mCitrine fluorescence/mKate2 fluorescence). Each dot represents the fluorescence value within a single cell. Thick horizontal bars represent the mean value, and error bars represent the 95% confidence interval as determined by bootstrapping, the values of which are shown in the **Table S4**. **(C)** *Stu1* MELT mutants disrupt Stu1-mCitrine localization to unattached kinetochores. *STU1-mCitrine* (wild type, SBY22493) and *stu1^2A^-mCitrine* (SBY23135) cells also containing fluorescent Spc110-mTurquoise2 and Mtw1-mKate2 were released from G1 into 15 µg/mL nocodazole for 1.5 hours to create unattached kinetochores and live imaged. The cells are displayed as in (A). **(D)** Quantification of mCitrine fluorescence from the experiment shown in (C). The same methods as (B) were used for quantification. **(E)** The degree of unattached kinetochore clustering was assessed using the bright kinetochore marker Mtw1-3xmYPet in *STU1-3xFLAG* (wild type, SBY23480), *stu1^T1034A^-3xFLAG* (SBY23772), *stu1^T1080A^-3xFLAG* (SBY23822), and *stu1^2A^-3xFLAG* (SBY23753) cells that were treated and imaged as in (A). Each genotype shows two representative cells. **(F)** Quantification of Mtw1-3xmYPet foci counts per cell from the experiment shown in (E). Dashed lines represent the mean foci count for each strain. *p-*values were generated using a Mann-Whitney U test. The wild type data in all graphs (*STU1-3xFLAG*, blue) are the same for easier comparison of the mutant distributions.

It was previously observed that, in addition to deleting TOG1, deleting Stu1’s CL region or inhibiting Mps1 kinase activity also disrupts Stu1 localization to unattached kinetochores^21,30^. We confirmed that removal of Stu1’s CL region (*stu1*^Δ^*^CL^*) prevents Stu1 localization to unattached kinetochores (**Figure S4**). We next tested whether preventing Stu1 MELT phosphorylation also affects Stu1 localization (**Figure 6C**). In the presence of nocodazole, Stu1^2A^-mCitrine levels on unattached kinetochores were drastically reduced compared to wild type controls (**Figure 6D**). These data suggest that Stu1 MELT phosphorylation is a key step in the recruitment of both Stu1 and Slk19 to unattached kinetochores.

We subsequently assessed the unattached kinetochore clustering phenotype during a nocodazole arrest in the *stu1* MELT mutants (**Figure 6E**). *Stu1^2A^* cells displayed more kinetochore foci compared to wild type controls, indicating a role for Stu1 MELT phosphorylation in clustering (2.72 foci per cell for wild type vs. 3.86 foci per cell for *stu1^2A^*, *p* < 0.001, **Figure 6F**). Interestingly, the single phospho-mutants did not show any kinetochore clustering defects. This may be explained by our findings that both single mutants can still recruit Slk19 to unattached kinetochores and co-purify with Stu1, albeit at lower levels. Collectively, these data indicate that phosphorylation of both Stu1 MELTs is required for maximal Slk19 recruitment to unattached kinetochores, but only one of the two MELTs (that supports reduced levels of Slk19) is sufficient to fully control clustering.

### Stu1 and Slk1G directly interact and form long, string-like filaments

We next asked whether kinetochore clustering might be mediated by a direct interaction between Stu1 and Slk19. To test this, we used an *in vitro* pull-down assay with purified components (**Figure 7A**). We recombinantly expressed and purified the Stu1 C-terminus (including the CL region and dimerization domain, hereafter referred to as Stu1-C), a recombinant Mps1 fragment, and native Slk19 (**Figure 7A, Figure S5A**). We immobilized Stu1-C on beads and performed phosphorylation by Mps1 or mock controls before adding soluble Slk19. As expected, we did not detect binding between Stu1-C and Slk19 in the absence of phosphorylation (**Figure 7A**). However, Stu1-C that was pre-incubated with ATP and recombinant Mps1 efficiently bound Slk19. Pre-incubation of Stu1-C with ATP alone failed to pull-down Slk19, ruling out any co-purifying kinase activity. Consistent with Mps1 phosphorylation of the Stu1-C fragment, Stu1-C exhibited a mobility shift during SDS-PAGE when incubated with active Mps1 and ATP (**Figure 7A**).

**Figure 7:**
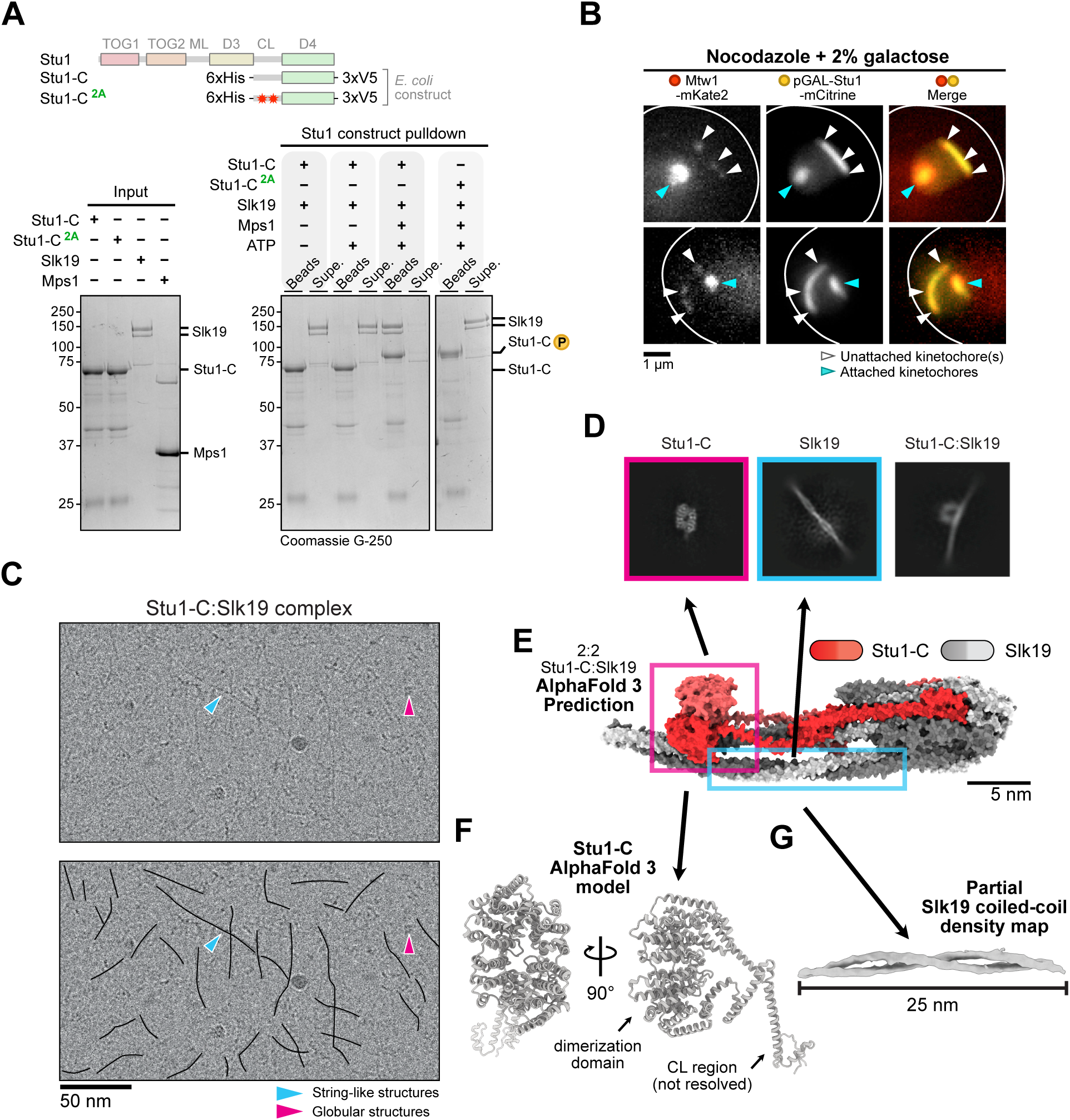
Stu1 and Slk1G directly interact and form long, string-like filaments. (A) Reconstitution of Stu1:Slk19 complex *in vitro* requires Mps1 phosphorylation of Stu1 MELTs. Top: schematic comparing recombinant Stu1 constructs to full-length Stu1. Red stars in Stu1-C^2A^ represent phospho-deficient alanine mutations of the MELT motifs. Bottom: Coomassie staining of SDS-PAGE gels of the pulldown experiment. The input (left) shows proteins prior to the pulldown (right). Note that in lanes indicating addition of recombinant Mps1 and/or ATP, Mps1 and/or ATP were added to beads pre-bound to Stu1-C or Stu1-C^2A^ and subsequently washed away before introducing Slk19. The supernatant lanes are unbound fractions after incubation with Slk19. **(B)** Overexpression of *STU1-mCitrine* during a nocodazole arrest drives Stu1 filament formation surrounding unattached kinetochores. *pGAL-STU1-mCitrine MTW1-mKate2* cells (SBY21509) were arrested in G1 (grown in 2% raffinose) and released into 15 µg/mL nocodazole to create unattached kinetochores and 2% galactose to induce overexpression. Cells were live imaged after 2 hours. **(C)** CryoEM micrograph of the Stu1-C:Slk19 reconstituted complex showing long, string-like filaments (blue arrowhead) and small, compact particles (pink arrowhead). Top: raw image; bottom: duplicate raw image with filaments highlighted with drawn lines. **(D)** 2D classifications of Stu1-C:Slk19 complexes using cryoEM. Left, pink: a diamond-shaped globular protein. Middle, blue: a coiled-coil protein. Right: the two proteins in complex. **(E)** AlphaFold 3 multimer model of a 2:2 Stu1-C:Slk19 phosphorylated complex. The pieces of the model that reflect the 2D classifications are boxed with their respective colors. **(F)** The diamond-shaped classification resembles the AlphaFold 3 model of the Stu1 dimerization domain (D4 domain). **(G)** A partial Slk19 coiled-coil density map at 7Å resolution spanning 25 nm in length.

We assessed whether Slk19 binding to Stu1-C depends on Stu1 MELT motif phosphorylation. We expressed and purified recombinant Stu1-C^2A^ in which both MELTs contained phosphodeficient alanine mutations (T1034A and T1080A) and repeated the binding assay. In alignment with our *in vivo* data, Slk19 failed to bind pre-phosphorylated Stu1-C^2A^ *in vitro* (**Figure 7A**). Stu1-C^2A^ exhibited a mobility shift like wild type during SDS-PAGE, indicating additional Mps1 phosphorylation sites on this fragment. Consistently, kinase assays with recombinant Mps1 and radioactive ATP-𝛾-^32^P on immobilized Stu1-C and Stu1-C^2A^ showed a reduction in phosphorylation on Stu1-C^2A^ (**Figure S5B**). These data align with our mass spectrometry analysis showing additional putative Mps1 sites in the CL region (**Figure 5C, Figure S2, Table S1**). Despite the unknown function of these additional sites, Mps1 phosphorylation of the two conserved Stu1 MELTs is required to directly recruit Slk19.

It has been proposed that Stu1 and Slk19 form an oligomeric network at unattached kinetochores because *STU1* overexpression leads to large structures surrounding unattached kinetochores^21, 29^. An oligomeric network may explain why preventing phosphorylation of Stu1 MELT motifs also disrupts Stu1 localization to unattached kinetochores, despite TOG1 being its kinetochore binding domain. We confirmed the overexpression phenotype of *STU1-mCitrine* cells using a galactose-inducible promoter during a nocodazole arrest (*pGAL-STU1-mCitrine*, **Figure 7B**). In these cells, unattached kinetochores appeared to spread out and decluster along the length of a large (> 1.5 μm) Stu1 filament, suggesting an oligomeric network may link unattached kinetochores together to facilitate clustering. We therefore set out to uncover structural information about the complex.

To test whether Stu1:Slk19 may form an oligomeric network, we subjected the reconstituted Stu1-C:Slk19 complex to cryogenic electron microscopy (cryoEM). In raw cryoEM micrographs, we observed many string-like filaments that spanned 100 nm in length (**Figure 7C**, blue arrow) and small compact particles (**Figure 7C**, pink arrow). Two-dimensional (2D) classifications show three structures: (1) a globular particle with two-fold symmetric density (**Figure 7D**, left), (2) a coiled-coil protein (**Figure 7D**, middle), and (3) both particles in complex (**Figure 7D**, right). To identify the particles, we used AlphaFold 3^47^ to model the structure of the Stu1-C:Slk19 complex. Although the stoichiometry of a Stu1:Slk19 complex is unknown, Slk19 has been reported to be a tetramer^48^ and Stu1, a dimer^30^. Unfortunately, a complex with this stoichiometry was too large to model using AlphaFold 3, even with the truncated Stu1-C construct. However, AlphaFold 3 models of a 2:2 Stu1-C:Slk19 (with phosphorylation included) predict Slk19 to be an elongated coiled-coil protein and Stu1-C to form a compact dimer with an interweaving of the phosphorylated CL region along the length of Slk19 (**Figure 7E**). We therefore conclude the globular particle is Stu1’s dimerization domain (**Figure 7F**) and the coiled-coil protein is Slk19.

We were unable to solve the structure of the complex due to severe preferred orientation and complex flexibility, preventing us from pinpointing the true position of the Stu1 MELT motif-Slk19 contact site. However, we were able to obtain a 7Å density map showing a partial, 25 nm-long Slk19 coiled-coil structure from 3D classifications (**Figure 7G**). Together, our work shows that Slk19 exists as an elongated coiled-coil and suggests that Stu1:Slk19 complexes form long, string-like filaments that could promote clustering of unattached kinetochores.

## DISCUSSION

Our work provides mechanistic insight into the molecular events that control unattached kinetochore clustering in budding yeast. We found that Mps1 kinase activity drives Stu1:Slk19 complex formation by phosphorylating Stu1’s MELT motifs to cluster unattached kinetochores. As unattached kinetochore clustering has been implicated in timely kinetochore capture^20^, we propose that Mps1 actively promotes initial kinetochore capture through Stu1:Slk19. Although we only probed its clustering function, this mechanism driven by Mps1 likely extends to Stu1 and Slk19’s other interdependent roles at unattached kinetochores during capture: promoting microtubule growth into the nuclear space to “search” for kinetochores, as well as stabilizing the capturing microtubule to prevent kinetochore detachment^21^.

It is intriguing that kinetochore clustering is nonessential during mitosis. Except for the *stu1* TOG1 point mutants showing a slight benomyl resistance, we did not find any growth phenotypes associated with disrupting unattached kinetochore clustering in *stu1^2A^* or *slk1S*Δ, suggesting the benomyl resistance is unrelated to Stu1’s clustering function. It was reported that *slk1S*Δ cells have a slight benomyl sensitivity^20^, a discrepancy that may be due to differences in genetic background. Nocodazole washout experiments of *slk1S*Δ cells show a notable delay in anaphase entry and in the completion of biorientation^20^, phenotypes that are likely to extend to *stu1^2A^* and *stu1* TOG1 mutants. As initial kinetochore attachment happens rapidly and efficiently during early S phase^49^, we speculate that there is sufficient time during the remainder of S phase to capture and biorient the kinetochores in the absence of Stu1:Slk19. It is also plausible that other proteins involved in kinetochore capture and transport such as Kar3 and Dam1^4, 50^ act to enhance the capture process in parallel with Stu1:Slk19, and that loss of only kinetochore clustering is not sufficient to cause delays. Lastly, although *SLK19* is nonessential during mitosis, it is essential for the completion of meiosis^51, 52^. Whether this is related to kinetochore clustering is unclear. It will be important to test if Stu1’s MELT motifs are required for the completion of meiosis in the future.

We uncovered the interplay between Stu1’s TOG1 domain and its disordered CL region, both of which are important for clustering. We discovered a basic patch on the surface of Stu1’s TOG1 domain required for its kinetochore localization. Although the kinetochore receptor of Stu1’s TOG1 domain and how its recruitment is regulated remain elusive, we favor the idea that this interaction is phospho-regulated similarly to the recruitment of Slk19 and spindle checkpoint proteins. It is likely that the only function of TOG1 in clustering is to recruit Stu1 to the kinetochore because restoring Stu1^ΔK94^ localization to kinetochores by fusing it to Bub3 rescues clustering defects. These rescue results also suggest Stu1’s receptor is likely positioned at the outer kinetochore near the Spc105 phospho-domain. It was previously reported that Mps1 is required for Stu1 localization to unattached kinetochores, suggesting Mps1 directly controls Stu1 recruitment^21^. However, since Stu1 MELT phosphorylation is required to visualize its co-localization with unattached kinetochores, it is unclear whether the role of Mps1 in Stu1 localization is to phosphorylate its receptor, the Stu1 MELTs, or both. It was also reported that Stu1 fails to localize to unattached kinetochores in *spc105-CA* cells in which Spc105 MELT motifs are unable to be phosphorylated^21^. Since the checkpoint proteins are not required for Stu1 recruitment^21, 29^, these findings suggest Stu1 may directly bind to Spc105 MELT motifs. However, given our results that Slk19 can bind phosphorylated MELTs and Slk19 is also required for Stu1 localization^21^, Slk19 binding to Spc105 MELTs is a more conceivable explanation. It will be interesting to investigate the role of the Spc105 MELTs in Stu1 and Slk19 localization in the future.

We discovered that Slk19 directly binds Stu1’s MELT motifs to control kinetochore clustering, similarly to the interaction of Bub3 with the phosphorylated MELTs of Spc105 to control the spindle checkpoint^35^. However, AlphaFold 3 models and our structural data suggest that Slk19 is an elongated coiled-coil protein, unlike the globular, 7-bladed β-propeller structure of Bub3^35^. Since we were unable to solve a high-resolution structure of the Stu1-C:Slk19 complex, it is unclear how Slk19 recognizes phosphorylated MELTs. Additional experiments will be required to determine whether the same Slk19 molecule binds both Stu1 MELTs simultaneously or whether separate Slk19 molecules interact with each MELT individually. However, *stu1^T1034A^* cells have a larger effect on disrupting Slk19-mCitrine recruitment to unattached kinetochores than *stu1^T1080A^* cells, consistent with a model where both MELTs are involved in recognition of the same Slk19 molecule with each site having a different contribution to Slk19 binding affinity. Regardless, only one MELT is sufficient for unattached kinetochore clustering, suggesting wild type amounts of Stu1:Slk19 at unattached kinetochores are well above the threshold required for clustering. Overall, these findings highlight the central importance of phosphorylated MELT motifs in unattached kinetochore regulation.

Our structural data revealed that Stu1-C:Slk19 complexes consist of long, string-like filaments (some > 100 nm in length), and we have confirmed that Stu1 overexpression forms a large filamentous structure that appears to decluster unattached kinetochores along its axis. These data fit with the previously proposed model that Stu1 and Slk19 may form an oligomeric network at unattached kinetochores^21, 29^. However, is unclear whether each filament we visualized using cryoEM is a repeating array of Stu1:Slk19 complexes or a single Stu1:Slk19 complex. If oligomers are indeed formed, Mps1-mediated oligomerization of Stu1:Slk19 filaments is reminiscent of the fibrous corona of metazoan kinetochores. The RZZ complex, along with the dynein-dynactin adaptor Spindly, oligomerize at unattached kinetochores in an Mps1-dependent manner at the outer kinetochore^13, 53, 54^. This scaffold recruits microtubule motors CENP-E and dynein involved in initial kinetochore capture and chromosome congression^11, 55–60^, as well as MAD1:MAD2 to control spindle checkpoint signaling^61^. Despite shared roles with the corona in the initial capture process, Stu1 and Slk19 do not control spindle checkpoint signaling. Additionally, the corona is built on the kinetochore of a single chromosome and has not been reported to cluster chromosomes, whereas Stu1 and Slk19 appear to “glue” multiple chromosomes together at their kinetochores^20, 62^. In the future, it will be interesting to determine whether Stu1 and Slk19 act alone to mediate clustering and capture, or whether this pathway is more corona-like in recruiting additional proteins for these processes.

In summary, our data suggest the following model for unattached kinetochore clustering (**Figure 8**). First, TOG1 binds unattached kinetochores through its basic patch to an unknown kinetochore receptor (**Figure 8A**). This initial binding event positions Stu1 in a favorable manner to kinetochore-localized Mps1, which phosphorylates Stu1’s MELT motifs to directly recruit Slk19 (**Figure 8A**). Phosphorylation of the MELTs is required to visualize both Stu1 and Slk19 on unattached kinetochores using microscopy assays, suggesting a positive feedback mechanism on Stu1 recruitment. This could be explained by the proposed “oligomerization model” where MELT phosphorylation initiates the oligomerization process by scaffolding alternating Stu1:Slk19 arrays^21^, which may be achieved through additional MELT-binding interfaces on the Slk19 tetramer. Alternatively, if oligomers do not form, an “allosteric model” is possible in which Slk19 binding to Stu1 MELTs further stabilizes TOG1 binding to the kinetochore (**Figure 8A**). Finally, the Stu1:Slk19 string-like filaments stimulate the clustering of unattached kinetochores (**Figure 8A**). How this works—and whether Stu1:Slk19 filament formation is sufficient for kinetochore clustering or whether other proteins are required—is unknown, but we speculate that filaments from individual unattached kinetochores may jointly coordinate their entrapment into a single functional unit (**Figure 8B**). Together, this work has identified another pathway that Mps1 controls specifically at unattached kinetochores to regulate chromosome segregation.

**Figure 8:**
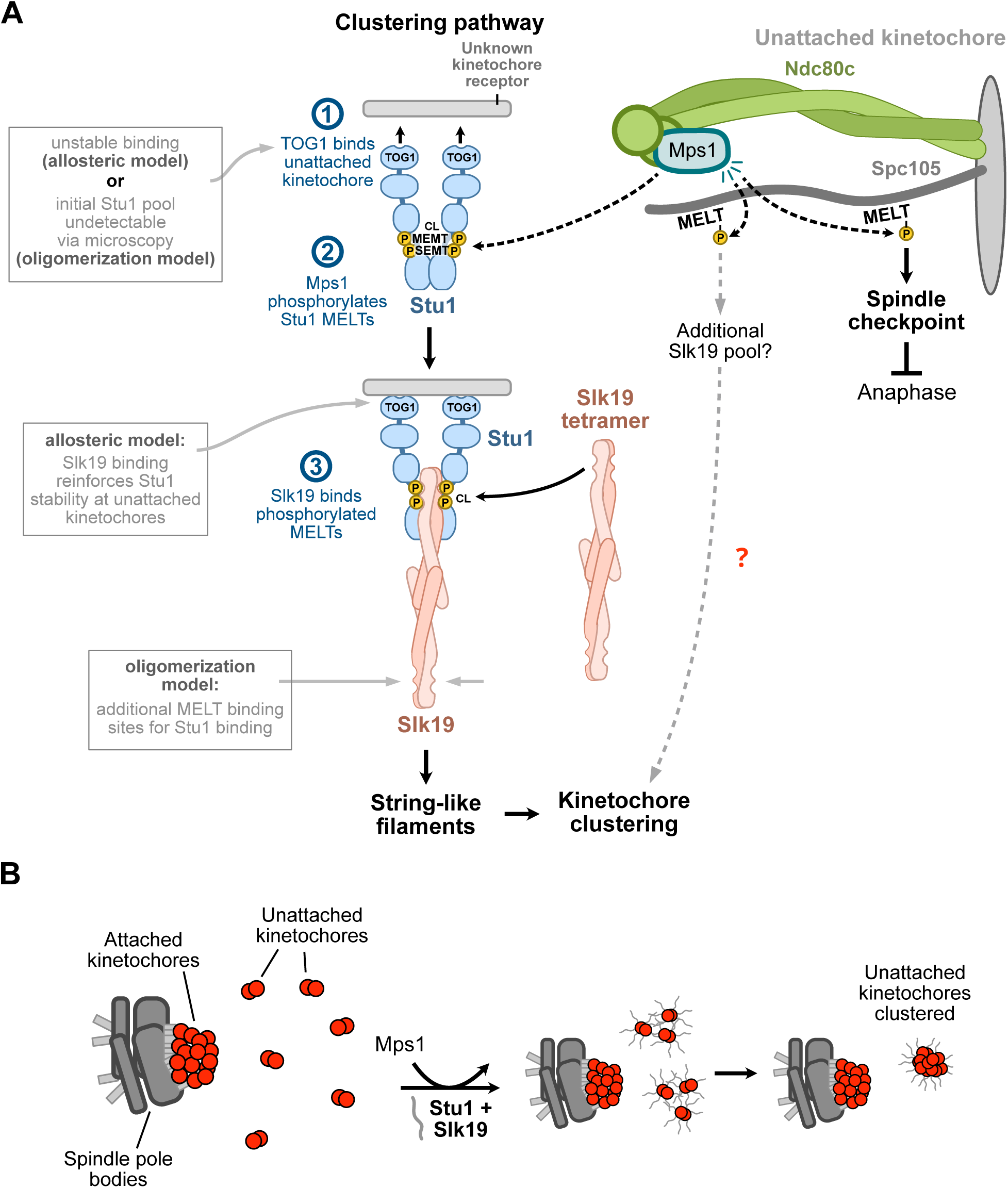
Model for the unattached kinetochore clustering pathway. (A) Mps1 activity at unattached kinetochores has separate roles in the initiation of the spindle checkpoint and kinetochore clustering. We propose the following model for unattached kinetochores to be fully competent in clustering: (1) Stu1 first binds unattached kinetochores via its TOG1 domain through its basic patch, bringing its CL region near kinetochore-bound Mps1. (2) Mps1 phosphorylates Stu1 MELT motifs within the CL region. (3) Slk19 binds to Stu1 MELTs and reinforces Stu1’s recruitment to the kinetochore, either through oligomerization or allostery. In the oligomerization model, additional MELT binding sites on the Slk19 tetramer may recruit additional Stu1. Although the role of Spc105 MELTs is unclear, we suggest Slk19, rather than Stu1, may be able to bind these MELTs and contribute to clustering. **(B)** Stu1:Slk19 filament formation at unattached kinetochores by Mps1 promotes the clustering of unattached kinetochores, possibly through a coordinated entanglement of the filaments between different unattached kinetochores.

## MATERIALS AND METHODS

### Yeast strain construction

All yeast strains used in this study are derivatives of the *Saccharomyces cerevisiae* W303 background (SBY3) and are listed in **Table S2**. Yeast strains were constructed using established genetic protocols^63^. Briefly, to generate gene products encoding C-terminal tags at endogenous loci, PCR products containing homologous ends to the directed locus were transformed^64^ (all plasmids used as templates are listed in **Table S3**). To generate *MTW1-3xmYPet*, integration vector pSB3562 was linearized within *MTW1* using SnaBI and transformed. To integrate ectopic constructs at *LEU2*, constructs were cloned into a *LEU2* integration vector and linearized with PmeI prior to transformation (see plasmid construction). For CRISPR editing, a plasmid expressing *S. pyogenes* Cas9 and a sgRNA directed to the edit site was co-transformed with a linearized repair template (either single-stranded or double-stranded)^65^ and selected using a *URA3* marker. Recovered colonies were plated on 5-FOA media to counter-select *URA3*^66^ and terminate Cas9 expression. Successful edits were verified by sequencing. Yeast containing the *mps1-1* allele and the *cdc5-1* allele were kind gifts from Andrew Murray (Harvard University, Cambridge, MA) and David Morgan (UCSF, San Francisco, CA), respectively.

### Plasmid construction

All plasmids are listed in **Table S3**. All sequences were verified using sequencing. To generate the *MTW1-3xmYPet* integration plasmid (pSB3562), pSB3554 (Addgene #168056) was linearized using PCR with primers SB8584 and SB8585 upstream of YPet start codon. This was assembled via the Gibson method^67^ with gBlock SB8586. The resulting plasmid (pSB3561) was linearized with BamHI and assembled via the Gibson method with MTW1 sequence (from gDNA using primers SB8599 and SB8600) to generate pSB3562.

To generate *pGAL-SLK1S-3xFLAG LEU2* integration plasmid (pSB3515), pSB2223 (pL300, a gift from Karsten Weis, ETH Zürich, Zürich, CH) was digested by PspOMI and XhoI. *pGAL-STU2* was amplified from yeast gDNA (SBY12342) using primers SB4373 and SB4374, digested by PsmsOMI and XhoI, and ligated to the backbone using T4 DNA ligase (M0202, New England Biolabs) into linearized pL300. This plasmid was linearized with PCR using SB8411 and SB8412 to remove *STU2* (backbone). *SLK1S-3xFLAG* was amplified from gDNA (SBY8964) using SB8409 and SB8410 and subsequently assembled via the Gibson method with the backbone.

To generate a *pSTU1-STU1-NLS-mCitrine-3xFLAG LEU2* integration plasmid (pSB3379), we first generated a *pSTU1-STU1-3xFLAG LEU2* integration plasmid (pSB3363) by amplifying pSB2223 (pL300) using primers SB7558 and SB7559 (backbone). *pSTU1-STU1-3xFLAG* was amplified from gDNA (SBY21199) using primers SB7555 and SB7557 and subsequently assembled via the Gibson method with the backbone. This plasmid (pSB3363) was amplified using primers SB7601 and SB7602 (backbone), and a fragment containing the SV40 nuclear localization signal (NLS) to ensure nuclear localization followed by mCitrine was amplified from pSB2936 using primers SB7611 and SB7612. The two resulting fragments were assembled via the Gibson method.

To generate *pSTU1-stu1*𝛥*TOG1-NLS-mCitrine-3xFLAG* (pSB3390) and *pSTU1-stu1*𝛥*CL-NLS-mCitrine-3xFLAG* (pSB3417) *LEU2* integration plasmids, parent plasmid pSB3397 was amplified using primers SB7642 and SB7643 (removing TOG1, residues 2-260) and primers SB7831 and SB7832 (removing CL, residues 997-1180), respectively, and the PCR products of each were treated with KLD enzyme mix (M0554, New England Biolabs).

To generate *pSTU1-stu1-T1034A-T1080A-NLS-mCitrine-3xFLAG LEU2* integration plasmid (pSB3518), we mutated each site successively using a Q5 site-directed mutagenesis kit (E0554, New England Biolabs). First, parent plasmid pSB3379 (*pSTU1-STU1-NLS-mCitrine-3xFLAG*) was mutagenized using SB8243 and SB8244 to introduce Thr1034Ala. This plasmid was then mutagenized using SB8440 and SB8441 to introduce Thr1080Ala.

To generate the CRSIPR-targeting plasmids (pSB3483, pSB3419, pSB3465), parent plasmid pSB3218 (pPGK1-SpCas9 GFP-sgRNA, a gift from Elçin Ünal, UC Berkeley, Berkeley, CA) was linearized using BsmBI and ligated with annealed protospacer oligonucleotides containing sticky-ends using instant-sticky end ligase master mix (M0370, New England Biolabs).

To generate 6xHis-Stu1-C-3xV5 (pSB3560) and 6xHis-Stu1-C-2A-3xV5 (pSB3564) recombinant expression plasmids, we first generated *STU1-3xV5-CxHis* in a pET-21b(+) (Novagen) backbone by amplifying *STU1-3xV5* from gDNA of strain SBY12019 using primers SB2967 and SB7674. The PCR product was purified on a column and re-amplified using SB8430 and SB8431 to add the 6xHis tag and assembled via the Gibson method into pET-21b(+) backbone generated by PCR using SB8438 and SB8439. The resulting plasmid (pSB3519) was amplified using SB8462 and SB8463 and treated with KLD enzyme mix (M0554, New England Biolabs) to remove all domains from *STU1* except CL and D4 resulting in plasmid pSB3524. For biochemistry purposes, the sequence encoding the 6xHis tag was removed from the sequence encoding the C-terminal end of *STU1* by PCR using SB8536 and SB8537, treated with KLD enzyme mix to generate pSB3559, re-amplified using SB8538 and SB7961, and treated with KLD enzyme mix to generate an N-terminal 6xHis tag resulting in pSB3560. To generate 6xHis-Stu1-C-2A-3xV5, we amplified pSB3560 using SB8601 and SB8602, omitting sequence encoding Stu1 MELTs, and amplified *stu1^2A^* sequence from pSB3518 using SB8450 and SB7602. The two fragments were assembled via the Gibson method to generate pSB3564.

To generate the 6xHis-Mps1(428-End) recombinant expression plasmid (pSB3601), full-length MPS1 was cloned into pSB3587 (pCDFDuet1 containing Lambda protein phosphatase downstream of the second RBS, a kind gift from Arshad Desai, UC San Diego, La Jolla, CA). pSB3587 was linearized with BamHI downstream of the first RBS and full-length *MPS1* was amplified using SB8727 and SB8728 from wild type gDNA containing homology to the backbone ends. The two fragments were assembled via the Gibson method to generate pSB3593. To remove the disordered N-terminus of Mps1, this plasmid was amplified using SB8767 and SB8768 and treated with KLD enzyme mix (New England Biolabs), resulting in a plasmid to co-express 6xHis-Mps1(428-End) and Lamba phosphatase (pSB3601).

To generate the *pGAL-STU1-mCitrine* integration plasmid (pSB3376), a *LEU2* integration plasmid containing the *pGAL1-10* promoter was amplified using primers SB7603 and SB7604. *STU1-mCitrine* was amplified from gDNA of SBY20768 using primers SB7605 and SB7606. The two PCR fragments were assembled via the Gibson method, resulting in pSB3376.

### Live cell fluorescence microscopy and quantification

For live cell imaging, log-phase cultures were grown in synthetic complete (SC) media (supplemented with 2% glucose or 2% raffinose for *pGAL* experiments, and 0.06% adenine), diluted to OD_600_ = 0.4, and arrested in G1 with 1 µg/mL α-factor for 3 hours (for auxin experiments, 2 mM indole-3-acetic acid (IAA) was added or an equivalent volume of DMSO as a control 30 minutes prior to harvest). Cells were harvested at 3,000 x g for 3 min and washed with yeast extract peptone (YEP) (supplemented with 2% glucose or 2% raffinose for *pGAL* experiments, and 0.02% adenine, 2 mM IAA or DMSO as a control for auxin experiments) for a total of 3 times. Cells were resuspended in YEP + 2% glucose + 0.02% adenine, supplemented with 1% DMSO, 15 µg/mL nocodazole, and 2 mM IAA or DMSO as a control (only for auxin experiments) to generate unattached kinetochores. For *pGAL* experiments, the same media was used except 2% galactose instead of glucose was added. For *mps1-1* experiments, resuspended cultures were shifted to 37 °C, harvested, and imaged after small buds formed. For imaging unperturbed metaphase cells, nocodazole was omitted, and cells were imaged after small buds formed. For all other experiments, resuspended cultures were harvested and imaged after 1.5-2 hours. To harvest, 1 mL of cells were centrifuged at 3,000 x g for 2 min and resuspended in 50 µL SC media supplemented with 2% glucose (or 2% galactose for pGAL experiments), 0.06% adenine, 1% DMSO, 1% peptone, and 15 µg/mL nocodazole (2 mM IAA or equivalent volume of DMSO was added for auxin experiments). 1.5 µL of cells were loaded onto a 1.5% agarose pad (dissolved in SC media with 2% glucose or galactose and 0.06% adenine) assembled onto a microscope slide and immediately imaged on a DeltaVision Ultra microscope with a 100x/1.4 PlanApo N oil-immersion objective (Olympus) using a 16-bit scientific complementary metal-oxide-semiconductor detector at 22 °C. Cells were imaged using the CFP-YFP-mCh filter set with excitation, emission as follows: CFP (438 nm, 475 nm), YFP (513 nm, 548 nm) and mCherry (575 nm, 625 nm) to image mTurquoise2, mCitrine or mYPet, and mKate2 signals, respectively at 100% power and 0.1 second exposure. DIC images were obtained using 32% power and 0.05 second exposure. Z-stacks of cells were obtained at 0.5 µm sections.

For kinetochore clustering quantification, cells containing the bright kinetochore marker Mtw1-3xmYPet were used due to a difficulty in visualizing individual kinetochore foci in Mtw1-mKate2 cells due to a low signal-to-noise ratio. These cells also contained Spc110-mTurquoise2 as a spindle pole body marker to verify the presence of collapsed poles as a readout for spindle disassembly by nocodazole prior to quantification. Strain identification was blinded and distinct kinetochore foci in the X, Y, and Z planes were counted manually per cell. The datasets were imported into RStudio, and histograms were generated using ggplot2. The Mann-Whitney U test was used to statistically compare distributions and generate *p*-values.

For fluorescence intensity measurements, intensity was measured using custom ImageJ macro scripts. Raw images of single cells were converted to maximum-intensity projections and the mean gray value of each channel at each kinetochore focus as well as the background fluorescence was quantified. Strain identification labels were blinded. The user manually thresholded the CFP (Spc110) and mCherry (Mtw1) channels for ROI selection. The macro distinguishes attached versus unattached kinetochores based on the co-localization of the kinetochore puncta with Spc110-mTurquoise2. The background signal of each channel was subtracted from the signal measured at each ROI (background-corrected mean gray value). The corrected Stu1-mCitrine or Slk19-mCitrine value at each kinetochore was divided by the Mtw1-mKate2 signal at that kinetochore (“normalized”). Each cell contained one normalized mCitrine measurement at attached kinetochores, with one or more normalized measurements at unattached kinetochores. In the case of more than one unattached kinetochore measurement, we calculated the mean of all normalized unattached kinetochore measurements to report a single unattached kinetochore measurement per cell. The datasets were imported into RStudio, and dot plots were created using ggplot2^68^. The dot plots display individual normalized values. The thick horizontal bars represent the mean value, and error bars represent the 95% confidence interval calculated by bootstrapping using ggplot2’s stat_summary() function. The statistical values are reported in **Table S4**. Each live imaging experiment was performed with three biological and technical replicates.

### Spindle checkpoint monitoring assay

Log-phase cultures grown in YEP supplemented with 2% glucose and 0.02% adenine (“media”) were diluted to OD_600_ = 0.4 into 25 mL and arrested in G1 with 1 µg/mL α-factor for 3 hours. Cells were centrifuged at 3,000 x g for 3 min, washed with media a total of 3 times, and resuspended in 25 mL media. At each indicated time point, the OD_600_ was measured, and 1 mL of cells were harvested at 3,000 x g for 2 minutes and cell pellets were immediately frozen in liquid nitrogen. The next day, Laemmli sample buffer (volume normalized to OD_600_) was added to cell pellets and lysed using 0.5 mm glass beads in a cell homogenizer (BioSpec Products). Samples were thoroughly boiled before SDS-PAGE followed by immunoblotting.

### Serial dilution growth assay

Yeast cultures were grown to saturation for 24 hours in 2 mL YEP supplemented with 2% glucose and 0.02% adenine. Saturated cultures were serially diluted 5-fold using sterile water into 6 columns of a 96-well plate and plated onto the indicated plates. Plates were grown at the indicated conditions for 1-3 days and imaged after sufficient growth.

### Sequence alignments

*STU1* homolog sequences of other fungi species were obtained using NCBI BLAST and aligned using the MUSCLE algorithm in the SnapGene software (www.snapgene.com).

### AlphaFold modeling

For structure prediction only involving Stu1 (and not Slk19), the Stu1 structure in the AlphaFold 2 database was used^32^. For multimer structure prediction involving both Stu1-C (residues 997 to 1513) and Slk19, AlphaFold 3^47^ was used with 2:2 stoichiometry and the models included phosphorylation sites detected across all mass spectrometry samples for both proteins (**Table S1**). PyMOL (The PyMOL Molecular Graphics System, Version 3.0 Schrödinger, LLC.) and UCSF ChimeraX^69^ were used to display and manipulate structures. For surface charge analysis, the APBS electrostatics plugin was used and visualized in PyMOL^70^.

### Protein biochemistry

#### Immunoprecipitation

500 mL or 1 L of log-phase cultures were grown to OD_600_ between 3 and 4 in YEP + 2% glucose + 0.02% adenine and harvested. For benomyl arrests, 500 mL of 2X benomyl media (YEP, 2% glucose, 0.02% adenine and 60 µg/mL methyl 1-[butylcarbamoyl]-2-benzimidazolecarbamate) was added to 500 mL log-phase cultures (30 µg/mL benomyl final concentration) when they reached OD_600_ between 3.0 to 4.0 and allowed to grow for 3 hours prior to harvest. Cells were harvested by centrifugation at 5,000 x g for 5 min. Pelleted cells were washed in 35 mL cold water containing 0.2 mM PMSF, pelleted at 3,000 x g for 3 min, and washed in 25 mL lysis buffer (Buffer H 0.15): 25 mM HEPES pH 8, 2 mM MgCl2, 0.1 mM EDTA, 0.5 mM EGTA, 15% glycerol, 0.1% NP-40, 150 mM KCl supplemented with phosphatase inhibitors (1 mM sodium pyrophosphate, 2 mM sodium β-glycerophosphate, 0.1 mM sodium orthovanadate, 5 mM sodium fluoride, and 0.2 µM microcystin-LR) and protease inhibitors (0.2 mM PMSF, 20 µg/mL leupeptin, 20 µg/mL pepstatin A, 20 µg/ml chymostatin). Cells were pelleted at 3,000 x g for 3 min and resuspended in lysis buffer using the following calculation: OD x mL culture x 1.25. Resuspended cells were frozen in liquid nitrogen and lysed using a Freezer-Mill (SPEX SamplePrep) and then ultracentrifuged at 98,000 x g for 90 min (for 1 L cultures) or 13,200 x g in a table-top centrifuge (for 500 mL cultures) at 4 °C. Protein concentration was measured using a Pierce BCA Protein Assay Kit (Thermo Scientific) to normalize concentrations using lysis buffer prior to immunoprecipitation. Protein G Dynabeads (Invitrogen) were crosslinked to α-M3DK antibody (GenScript) that recognizes the 3xFLAG epitope tag using dimethyl pimelimidate (Sigma-Aldrich). Beads were added to lysates (15 µL beads per 12.6 mg protein) and incubated at 4 °C with rotational mixing for 3 hr. Beads were washed 3 times with lysis buffer containing 2 mM DTT, protease inhibitors, and phosphatase inhibitors, washed twice with the same buffer without DTT, and eluted in lysis buffer containing 0.5 mg/mL 3xFLAG peptide (Sigma-Aldrich) in half the bead volume by gentle agitation for 30 min at 23 °C. All immunoprecipitation experiments were performed on at least two biological and technical replicates.

#### Immunoblotting and silver stain analysis

All samples were boiled in Laemmli sample buffer and run on SDS-PAGE gels. For immunoblotting, proteins were transferred to a 0.45 µm nitrocellulose membrane using a wet apparatus (Hoefer). Membranes were blocked using 4% nonfat milk in PBST and incubated in primary antibody overnight, rotating at 4 °C. The following commercial primary antibodies were used: α-PGK1 (Invitrogen; 4592560; 1:10,000), α-M3DK (to recognize FLAG epitopes, GenScript; 1:10,000), α-Myc (Cell Signaling; 71D10; 1:1,000), and α-V5 (Invitrogen; R96025; 1:5,000). For custom antibodies: α-Ndc80 (OD4; 1:10,000) was a kind gift from Arshad Desai (UC San Diego, La Jolla, CA), and α-Spc105 (PAC4065, 1:10,000, as published in ^71^). Primary antibodies were washed away with PBST and blots were probed with secondary antibody with gentle rotation for 1 hr at 23 °C. Secondary antibodies used were sheep anti-mouse conjugated to HRP (Cytiva; NA931V; 1:10,000) and donkey anti-rabbit conjugated to HRP (Cytiva; NA934V; 1:10,000), detected using SuperSignal West Dura Chemiluminescent Substrate (ThermoFisher Scientific), and imaged using a ChemiDoc MP Imaging System (Bio-Rad). In immunoblots where two protein bands run at the same molecular weight, the membrane was probed for the first band as described above and then stripped by incubation in stripping buffer (2% SDS, 50 mM TRIS pH 6.8, and 100 mM beta-mercaptoethanol) on a rotator at 50 °C for 25 min, followed by re-blocking and incubation with the second primary antibody overnight. For silver stain analysis, samples were run on 4-12% Bis-Tris NuPAGE gels (Invitrogen) using MES buffer and stained using the Silver Quest Silver staining kit (Invitrogen).

#### Native and recombinant protein purification

Rosetta 2(DE3) pLysS cells (Novagen) were used for recombinant expression. All cultures were grown in Terrific Broth (12 g/L tryptone, 24 g/L yeast extract, 0.4% v/v glycerol, 0.017M KH_2_PO_4_, 0.072M K_2_HPO_4_) containing 34 µg/mL chloramphenicol, and 100 µg/mL ampicillin (for Stu1 constructs) or 50 µg/mL streptomycin (for Mps1). To express 6xHis-Stu1-C-3xV5 and 6xHis-Stu1-C-2A-3xV5, we transformed cells with pSB3560 and pSB3564, respectively. For Mps1(428-End), we transformed cells with pSB3601 to dually express 6xHis-Mps1(428-End) and Lambda phosphatase. In the morning, 1 L cultures were inoculated from 5 mL overnight cultures and allowed to grow to OD_600_ ∼0.8. Protein expression was induced with 0.1 mM IPTG for 16 hr at 18°C. Cells were pelleted at 5,000 x g for 5 min and resuspended at a ratio of 3 mL lysis buffer per g of pellet (for Stu1 constructs: 25 mM HEPES, pH 8, 2 mM MgCl_2_, 15% glycerol, 0.1% NP-40, 500 mM KCl, 0.5 mM TCEP; for Mps1 constructs: 50 mM HEPES pH 7.4, 500 mM NaCl, 5% glycerol, 0.5 mM TCEP) supplemented with protease inhibitors (40 µg/mL leupeptin, 40 µg/mL pepstatin A, 40 µg/ml chymostatin, and 1 mM PMSF) and 10 mM imidazole. Cells were lysed using sonication. The extract was spun 14,000 x g for 30 min to pellet debris, and clarified lysate was incubated with equilibrated HisPur cobalt resin (ThermoFisher Scientific) on rotational mixing for 1 hr. Resin was collected in a gravity column.

For Stu1 constructs, the column was washed with 10X bed volume lysis buffer supplemented with 20 mM imidazole, then washed with 10X bed volume lysis buffer (containing 150 mM KCl instead of 500 mM KCl) supplemented with 20 mM imidazole. Protein was eluted in 25 mM HEPES pH 8, 2 mM MgCl_2_, 15% glycerol, 0.1% NP-40, 150 mM KCl supplemented with protease inhibitors and 150 mM imidazole. Eluted protein was buffer exchanged into Buffer H 0.15 by immobilization on α-V5 beads immediately before pulldown experiments.

For Mps1, the column was washed with 10X bed volume lysis buffer supplemented with 20 mM imidazole, then washed with 10X bed volume lysis buffer (containing 150 mM NaCl instead of 500 mM NaCl) supplemented with 20 mM imidazole. Protein was eluted in 50 mM HEPES pH 7.4, 150 mM NaCl, 5% glycerol, and 1 mM TCEP supplemented with protease inhibitors and 150 mM imidazole. Eluted protein was buffer exchanged and concentrated via diafiltration using Amicon Ultra 4 (10K MWCO, EMD Millipore) into Buffer H 0.15 (omitting EGTA and EDTA) containing 0.2 mM PMSF and 1 mM TCEP and flash frozen in liquid nitrogen. The final concentration was estimated using SDS-PAGE followed by Coomassie G-250 staining comparing samples to albumin standards.

To purify native Slk19, we followed the immunoprecipitation protocol with the following modifications: 2 L of strain SBY23482 were grown in YEP with 2% raffinose and 0.02% adenine to OD_600_ ∼2.0, at which point galactose was added to 2% to induce overexpression and cells were shifted to 37 °C to inactivate Mps1. Harvested cells were resuspended in lysis buffer (Buffer H 0.75: 25 mM HEPES pH 8, 2 mM MgCl2, 0.1 mM EDTA, 0.5 mM EGTA, 15% glycerol, 0.1% NP-40, 750 mM KCl supplemented with phosphatase and protease inhibitors). After α-FLAG bead incubation, beads were washed 3x with Buffer H 1.0 (25 mM HEPES pH 8, 2 mM MgCl2, 0.1 mM EDTA, 0.5 mM EGTA, 15% glycerol, 0.1% NP-40, 1 M KCl) and 3x with Buffer H 0.15, both supplemented with phosphatase and protease inhibitors. Beads were eluted with 3x FLAG peptide (0.5 mg/mL) in half the bead volume, eluate collected, and re-eluted in fresh peptide. Both eluates were combined and concentrated using an Amicon Ultra 0.5 filter (10K MWCO) at 14,000 x g for 10 min. The final concentration was estimated using SDS-PAGE followed by Coomassie G-250 staining comparing samples to albumin standards.

#### Stu1-C Slk1G *in vitro* pulldown

Purified Stu1-C or Stu1-C^2A^ were immobilized on α-V5 conjugated Protein G Dynabeads (ThermoFisher Scientific) by incubation with column-eluted protein for 1.5 hr with rotational mixing at 4 °C. Beads were washed 2x with Buffer H 0.15 (omitting EGTA and EDTA) supplemented with 1 mM TCEP and 0.2 mM PMSF. Buffer was removed and beads were incubated with ATP Buffer 4 (25 mM HEPES pH 8, 10 mM MgCl2, 15% glycerol, 0.1% NP-40, 150 mM KCl, 1 mM TCEP, and 1 mM ATP) containing ∼1.2 µM recombinant Mps1 for 15 min at room temperature with occasional gentle mixing. As a negative control for phosphorylation, ATP was omitted from the buffer and its volume replaced with H_2_O. As a negative control for kinase activity, Mps1 was omitted. Beads were washed 2x with wash buffer (Buffer H 0.15) containing phosphatase inhibitors (1 mM sodium pyrophosphate, 2 mM sodium β-glycerophosphate, 0.1 mM sodium orthovanadate, 5 mM sodium fluoride, and 0.2 µM microcystin-LR) and protease inhibitors (0.2 mM PMSF, 20 µg/mL leupeptin, 20 µg/mL pepstatin A, 20 µg/ml chymostatin), buffer was removed, and samples were incubated with natively purified Slk19-3xFLAG (∼320 nM per reaction) for 15 min at room temperature with occasional gentle mixing. For each binding reaction, we estimated the concentration of immobilized Stu1-C and Stu1-C^2A^ to be roughly 600 nM using BSA standards. The unbound fraction was removed, saved, and resuspended and boiled in Laemmli sample buffer. Beads were washed 2x with wash buffer and boiled in Laemmli sample buffer to elute bound proteins. The bound and unbound fractions were analyzed using SDS-PAGE followed by Coomassie G-250 staining. Pulldown experiments were performed on at least two biological and technical replicates.

#### Radioactive kinase assays

0.6 mg (20 µL) α-V5 conjugated Protein G Dynabeads (Invitrogen) were incubated with purified Stu1-C or Stu1-C^2A^ for 1 hr at 4 °C with rotational mixing. Beads were washed a total of 3 times with Buffer H 0.15 (omitting EGTA and EDTA) supplemented with phosphatase inhibitors, protease inhibitors, and 1 mM DTT. After the last wash, buffer was removed and samples were treated with radioactive ATP buffer (50 mM Tris pH 7.4, 1 mM DTT, 25 mM sodium β-glycerophosphate, 5 mM MgCl_2_, 10 µM ATP, and 2.5 µCi ATP-𝛾-^32^P) containing ∼260 nM recombinant Mps1. Samples were incubated for 15 minutes at room temperature with occasional agitation and washed 2x with Buffer H 0.15 containing protease and phosphatase inhibitors. All buffer was removed, and samples were eluted in 20 µL Laemmli sample buffer with boiling. 5 µL of sample was run on a 4-12% Bis-Tris NuPAGE gel (Invitrogen), silver stained, and imaged. The gel was dehydrated using a vacuum gel-drying system (Hoefer) for 1 hr at 80 °C and exposed to a phosphor screen (Molecular Dynamics) for 10 min. The phosphor screen was imaged using an Amersham Typhoon (Cytiva) using default settings. This assay was performed with two technical replicates.

#### Phosphatase assay

Stu1-3xFLAG was immunoprecipitated as described above, except the final two wash steps omitted phosphatase inhibitors. After the last wash, the sample was split in half. The first sample was treated with Buffer H 0.15 supplemented with 1 mM MnCl_2_ and 120 units of Lambda Protein Phosphatase (P0753, New England Biolabs, Ipswich, MA, USA). The second sample was treated the same, except phosphatase inhibitors were added as a negative control (1 mM sodium pyrophosphate, 2 mM sodium β-glycerophosphate, 0.1 mM sodium orthovanadate, 5 mM sodium fluoride). Samples were thoroughly mixed and incubated at 30 °C for 20 min with occasional mixing. Supernatant was collected (unbound fraction), and beads were washed 2x with Buffer H 0.15 containing protease and phosphatase inhibitors before eluting in 0.5 mg/mL 3xFLAG peptide. Samples and unbound fractions were analyzed using SDS-PAGE and immunoblotting. The phosphatase assay was performed on two biological and technical replicates.

#### Mass spectrometry

Log-phase *STU1-3xFLAG cdc20-AID OsTIR1* cells (SBY20662) were grown to OD_600_ ∼3.0 (as for an immunoprecipitation experiment), cultures were split in 3 ways, and samples were treated with 0.1% DMSO (control, asynchronous), 30 µg/mL benomyl with 0.1% DMSO (metaphase arrest), or 1 mM auxin (indole-3-acetic acid) with 0.1% DMSO (metaphase arrest) for 2.5 hours at 23 °C prior to harvesting. Stu1-3xFLAG was immunoprecipitated as described above, with two additional washes in 50 mM Tris pH 8.3, 75 mM KCl, 1 mM EGTA, and eluted with 0.2% RapiGest (Waters Corporation, Milford, MA, USA) in 50 mM ammonium bicarbonate. Eluates were reduced with 10 mM DTT at 56 °C for 45 min and alkylated with 55 mM 2-chloroacetamide in the dark at room temperature for 30 min. Solutions were incubated with 250 ng Trypsin (Promega) overnight at 37 °C with mixing. Samples were allowed to equilibrate to ambient temperature, acidified with 30% formic acid, and shaken vigorously to degrade the RapiGest. Samples were spun hard for 5 min, transferred to a new low binding Eppendorf tube, and dried on a speedvac. The dried samples were resuspended in 30 µL 85% acetonitrile 15 mM ammonium formate, pH 2.8. The peptide solution was desalted over 1 cm TopTip packing material (Poly-LC-) packed in a 10 µL pipette tip. Material was washed 3x with 50 µL 85% acetonitrile 15 mM ammonium formate, pH 2.8, eluted with 50 µL 15 mM ammonium formate pH 2.8, and dried on a speedvac.

Desalted samples were resuspended in 2% acetonitrile in 0.1% formic acid and analyzed by LC/ESI MS/MS with an Easy-nLC 1000 (Thermo Scientific) nano HPLC system coupled to a tribrid Orbitrap Fusion mass spectrometer (Thermo Scientific). In-line de-salting was accomplished using a reversed-phase trap column (100 µm × 20 mm) packed with Magic C_18_AQ (5 µm 200Å resin; Michrom Bioresources, Bruker) followed by peptide separations on a reversed-phase column (75 μm × 270 mm) packed with Magic C_18_AQ (5 µm 100Å resin; Michrom Bioresources, Bruker) directly mounted on the electrospray ion source. A 150 min gradient from 5% to 30% acetonitrile in 0.1% formic acid at a flow rate of 300 nL/min was used for chromatographic separations. The heated capillary temperature was set to 300 °C and a static spray voltage of 2200 V was applied to the electrospray tip. The Orbitrap Fusion instrument was operated in the data-dependent mode, switching automatically between MS survey scans in the Orbitrap (AGC target value 500,000, resolution 120,000, and maximum injection time 50 ms) with MS/MS spectra acquisition in the linear ion trap using quadrupole isolation. A 3 second cycle time was selected between master full scans in the Fourier-transform (FT) and the ions selected for fragmentation in the HCD cell by higher-energy collisional dissociation with a normalized collision energy of 27%. Selected ions were dynamically excluded for 20 seconds and exclusion mass by mass width +/-10 ppm.

Data analysis was performed using Proteome Discoverer 2.5 (Thermo Scientific). The data were searched against SGDyeast (SGD July 28, 2002) and cRAP (http://www.thegpm.org/crap/) FASTA files. Trypsin was set as the enzyme with maximum missed cleavages set to 2. The precursor ion tolerance was set to 10 ppm, and the fragment ion tolerance was set to 0.6 Da. Variable modifications included oxidation on methionine (+15.995 Da), and phosphorylation on serine, threonine and tyrosine (+79.966 Da). Dynamic modification on the protein N-terminus included acetylation (+42.011 Da). Static modifications included carbamidomethyl on cysteine (+57.021 Da). Data were searched using Sequest HT. All search results were run through Percolator for scoring and identified peptides were filtered for 1% peptide-level false discovery rate using *q* value of 0.01.

#### Kinase prediction of phosphorylation sites

The Proteome Discoverer results were exported as Excel spreadsheets and parsed by Python scripts as previously described^72^ to extract and compile phosphorylation sites that contained a confidence score ≥ 75% and ≥ 2 peptide spectrum matches (PSMs) for each dataset to generate **Table S1**. To identify a putative kinase, the sequence surrounding each phosphorylation site was searched for kinase consensus motifs of Mps1, Ipl1, Cdk1, and Cdc5 using regular expressions (see **Table S5** for all expressions used). For Mps1 specifically, we used the regular expression `[DENCQ][A-Z]([ST])[^PN]`, where the position in parentheses is the phospho-acceptor^45, 46^. Since Mps1 sites can also be primed at the-2 position by another phospho-residue^73^, we also included the regular expression `[ST][A-Z]([ST])[^PN]`. For primed sites, the site was labeled an Mps1 site (in **Figure 5C, Figure S2, and Figure S3**) if the-2 position was detected in the dataset as being phosphorylated. We also determined whether each site had been previously reported on the Saccharomyces Genome Database (SGD) (**Table S1**), although we note that not all reported phosphorylation in the literature is annotated in the SGD.

### Structural Biology

#### Negative stain EM

Purified Stu1-C was immobilized on α-V5 conjugated Protein G Dynabeads using an identical protocol as the *in vitro* pulldown described earlier. Beads were incubated in ATP Buffer 4 with purified native Slk19-3xFLAG and recombinant Mps1 for 20 min with occasional gentle mixing at room temperature. Beads were washed 2x with Buffer H 0.15 (omitting NP-40 and glycerol) and the complex was eluted with 0.5 mg/mL V5 peptide (Proteintech) dissolved in the same buffer. Complexes and Slk19-3xFLAG alone were diluted and deposited on 400 mesh continuous carbon grid (Electron Microscopy Sciences) for 1 min. The grids were washed twice with MilliQ water and stained with 0.75% Uranyl Formate (Electron Microscopy Sciences) for 1 min. Grids were air dried before loading into Talos L120C (ThermoFisher Scientific). The data was collected at 1.991 Å/pixel and processed in cryoSPARC^74^.

#### CryoEM data collection

3 µL undiluted Stu1-C:Slk19 complex was applied on plasma cleaned 300 mesh, 2/1 holey carbon grid with 2 nm continuous amorphous carbon film (Electron Microscopy Sciences) at 4 °C with 100% relative humidity and vitrified using a Vitrobot Mark IV (ThermoFisher Scientific). The grids were stored in liquid nitrogen until usage. The Stu1-C:Slk19 complex was imaged under Titan Krios G3 transmission electron microscope (ThermoFisher Scientific) operated at 300 kV. The microscope is equipped with a Gatan K3 summit direct detection camera (Gatan). A total of 8,466 micrographs were collected at a pixel size of 1.088 Å. For each micrograph, a 50-frame movie stack was collected with total exposure at 50e/Å^2^. An in-column energy filter was used with a slit width of 8 eV.

## Data processing

The collected micrographs were motion-corrected and dose-weighted in cryoSPARC ^74^. Contrast transfer function (CTF)-corrected micrographs were used for blob picking, and particles that belonged to well-resolved 2D classes were used for topaz picking ^75^ for second round particle picking. The newly picked particles were used for multiple rounds of 2D classification. Since Stu1 and Slk19 had very different shapes and sizes, we averaged them separately. 149,220 particles that belong to Slk19 were used for *ab initio* reconstruction and 3D classification. 43,128 particles from well-resolved 3D class were used for NU-refinement and generated 6.8 Å density map. The overall resolution was assessed using the gold-standard criterion of Fourier shell correlation (FSC), with a cutoff at 0.143, between 2 half maps from 2 independent half-sets of data. 487,648 particles were assigned to be Stu1, of which we were only able to get good 2D classifications. Density maps with severe, preferred orientation were obtained and not displayed in the text.

## Supplemental Material

**Figure S1** shows additional characterization of Stu1’s TOG1 domain. **Figure S2** shows the conservation of Stu1 MELT motifs using a multiple sequence alignment of the CL region with other budding yeast species. **Figure S3** displays the mass spectrometry coverage map of Stu1-3xFLAG immunoprecipitation experiments. **Figure S4** shows deletion of Stu1’s CL region disrupts its localization to unattached kinetochores, similarly to *stu1^2A^*. **Figure S5** shows protein purification results prior to the pulldown experiment and a radioactive kinase assay of Stu1-C. **Table S1** shows a complete list of phosphorylation detected from mass spectrometry analysis of Stu1-3xFLAG and kinase predictions of each site. **Table S2** lists the yeast strains used in this study. **Table S3** lists the plasmids used in this study.

**Table S4** displays the statistical analysis outputs used in this study and organized by figure. **Table S5** shows the regular expressions used to predict kinases in the phosphorylation data. **Table S6** lists the oligonucleotides and gBlocks used in this study.

## Data availability statement

All reagents are available from the corresponding author upon request. Raw mass spectrometry data (data generated in **Table S1**) is available at ftp://massive-ftp.ucsd.edu/v10/MSV000099081/. Python code with all relevant files to generate **Table S1** is available at www.doi.org/10.5281/zenodo.18217341. The cryoEM density map of Slk19 is available from EM Databank (EMDB) with accession number EMD-75100.

## Acknowledgements

We thank members of the Biggins and Asbury labs for critical reading of the manuscript, and members of the Seattle Mitosis group for helpful feedback and discussions. We thank Phil Gafken and Lisa Jones of the Proteomics & Metabolomics Core facility at Fred Hutchinson Cancer Center for their expertise in processing and analyzing mass spectrometry samples. We thank Elçin Ünal, Arshad Desai, Kim Nasmyth, Adele Marston, David Morgan, Karsten Weis, and Andrew Murray for providing strains and reagents. This work was supported by NIH P30 CA015704 award to the Proteomics & Metabolomics Shared Resource of the Fred Hutch/University of Washington Cancer Consortium and by NIH grant NIGMS R35 GM149357 to SB who is also an investigator of the Howard Hughes Medical Institute.

## Author contributions

Conceptualization: DRM, MJ, SB. Data curation: DRM, MJ, GMM. Formal analysis: DRM, MJ. Funding acquisition: SB. Investigation: DRM, MJ, GMM. Methodology: DRM, MJ, SB. Project administration: SB. Resources: DRM, MJ, SB. Software: DRM. Supervision: SB. Validation: DRM, MJ. Writing – original draft: DRM, MJ, SB. Writing – reviewing & editing: DRM, MJ, GMM, SB.

## Notes

ftp://massive-ftp.ucsd.edu/v10/MSV000099081/

https://zenodo.org/records/18217341

https://www.ebi.ac.uk/emdb/EMD-75100

### Competing Interest Statement

The authors have declared no competing interest.

### Summary of Updates

We have performed additional experiments to analyze the Stu1 CL deletion mutant and have revised the text to be more clear.

