## Supplemental Data for "Kinetochore clustering is mediated by Mps1 phosphorylation of conserved MELT motifs in Stu1"

**SUPPLEMENTAL MATERIAL**

Supplemental Figures

Figure S1: Additional characterization of Stu1's TOG1 domain.
Figure S2: The MELT motifs within Stu1's CL region are conserved in *Saccharomycetes*.
Figure S3: Mass spectrometry coverage maps of Stu1-3xFLAG immunoprecipitates.
Figure S4: Deletion of Stu1's CL region disrupts its localization to unattached kinetochores.
Figure S5: Stu1 is a substrate of the Mps1 kinase *in vitro*.

Supplemental Tables

Table S1: Detected phosphorylation sites from Stu1-3xFLAG immunoprecipitation mass spectrometry experiments - *attached as Excel file*
Table S2: Yeast strains used in this study.
Table S3: Plasmids used in this study.
Table S4: Statistics of image quantification.
Table S5: Regular expressions used to predict kinases in phosphorylation data.
Table S6: Oligonucleotides and gBlocks used in this study.

### SUPPLEMENTAL FIGURES

#### Figure S1

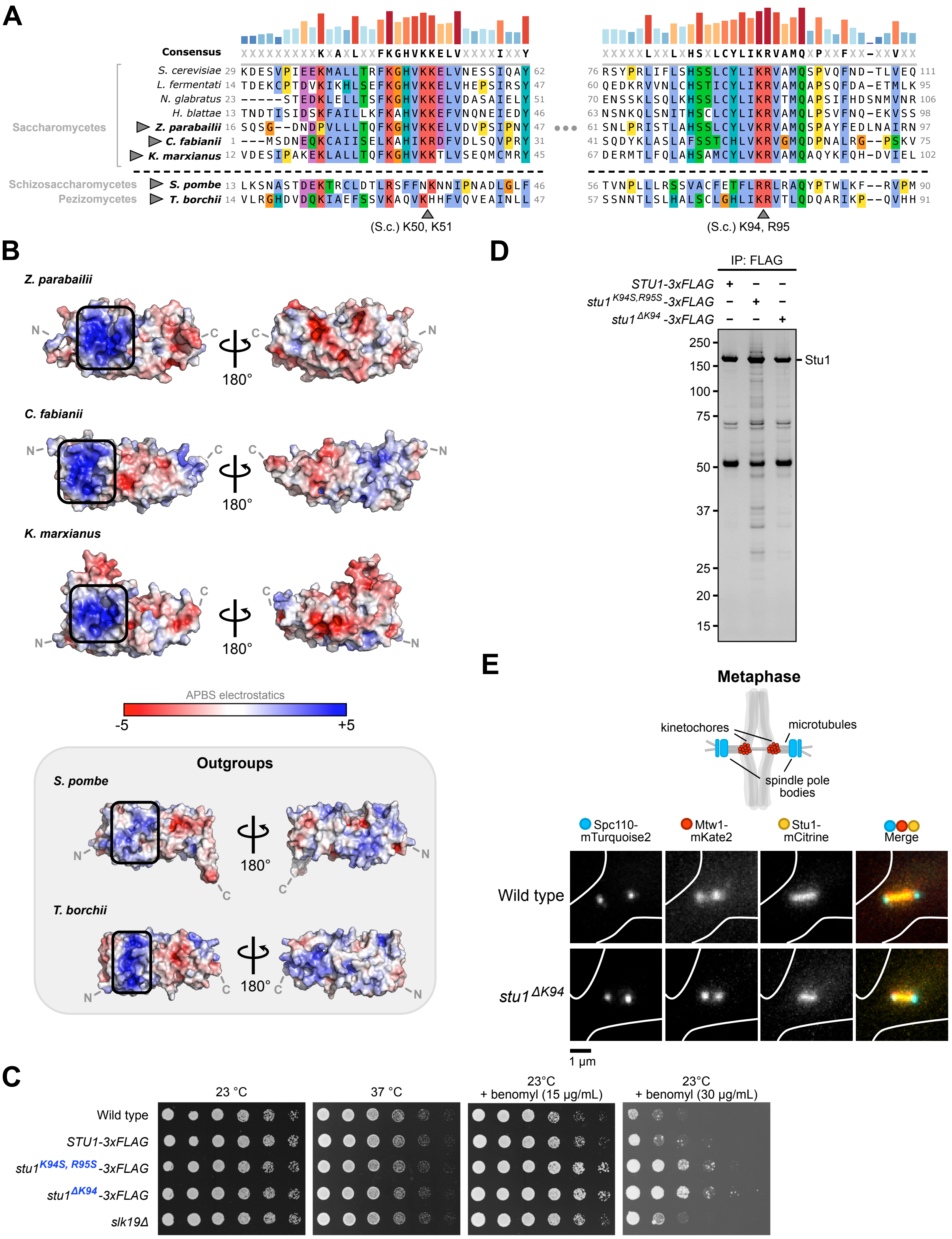

**Figure S1: Additional characterization of Stu1's TOG1 domain. (A)** Two snapshots of a multiple sequence alignment of Stu1's TOG1 domain from several yeasts belonging to the *Saccharomycetes* class (budding yeasts), as well as two fungi belonging to other clades (*S. pombe*, a fission yeast; and *T. borchii*, a truffle mushroom). The *S. cerevisiae* Stu1 residues K50 and K51 are also situated within the basic patch. **(B)** Surface electrostatic potential analysis of TOG1 AlphaFold models of select species within the alignment in (A). Note that TOG1 of yeasts belonging to the *Saccharomycetes* clade have a very defined basic patch like *S. cerevisiae* TOG1. The basic patch is less well-defined in the outgroups. **(C)** Stu1 TOG1 CRISPR mutants display an increased fitness on benomyl. Wild type (SBY3), *STU1-3xFLAG* (SBY21199), *stu1*$^{K94S, R95S}$*-3xFLAG* (SBY21941), *stu1*$^{\Delta K94}$*-3xFLAG* (SBY21937), and *slk19Δ* (SBY7513) cells were grown to saturation and five-fold serial dilutions were plated (from left to right), incubated at the indicated conditions for 24-72 hours, and imaged after sufficient growth. **(D)** The Stu1$^{K94S, R95S}$ mutant protein displays slightly enhanced degradation products compared to Stu1$^{\Delta K94}$. Benomyl-arrested lysates from *STU1-3xFLAG* (SBY21197), *stu1*$^{K94S, R95S}$*-3xFLAG* (SBY21810), and *stu1*$^{\Delta K94}$*-3xFLAG* (SBY21811) cells were immunoprecipitated using $\alpha$-FLAG beads and analyzed by silver staining. **(E)** Disrupting Stu1's basic patch does not alter its localization to the mitotic spindle. *STU1-mCitrine* (wild type, SBY21184) and *stu1*$^{\Delta K94}$*-mCitrine* (SBY22568) cells containing fluorescent Mtw1-mKate2 and Spc110-mTurquoise2 were released from a G1 arrest and live imaged after small buds appeared. A representative cell from each genotype is displayed as in Figure 1F.

Figure S2

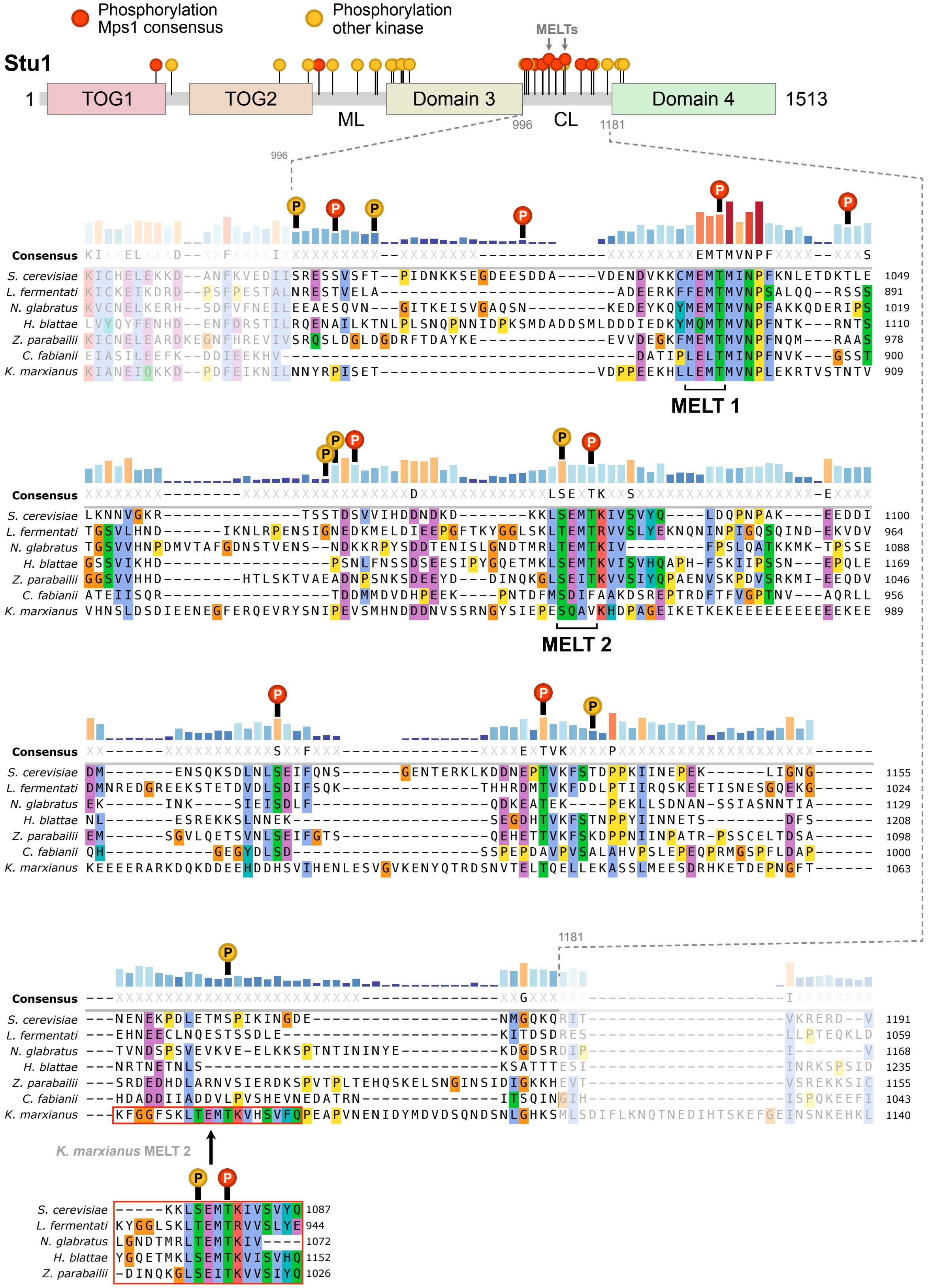
Phosphorylation Mps1 consensus
Phosphorylation other kinase
MELTs
Stu1
1
TOG1
TOG2
ML
Domain 3
996
CL
1181
Domain 4
1513
Consensus
S. cerevisiae
L. fermentati
N. glabratus
H. blattae
Z. parabailii
C. fabianii
K. marxianus
MELT 1
MELT 2
K. marxianus MELT 2

**Figure S2: The MELT motifs within Stu1's CL region are conserved in *Saccharomycetes*.** Multiple sequence alignment of Stu1's CL region from several yeasts belonging to the *Saccharomycetes* class (budding yeasts). Top: schematic of Stu1 domain organization (as displayed in **Figure 5C**). Bottom: protein alignment of the CL region with phosphorylation sites noted above the respective residue. The phosphorylation sites displayed are cumulatively detected phosphorylation sites from all conditions (asynchronous, benomyl-arrest, and metaphase-arrested; for individual conditions and coverage maps, see **Figure S3**). Red phosphorylation marks represent sites matching the Mps1 consensus. Much of the CL region contains large stretches of residues that lack conservation. However, the MELT motifs are conserved. The bars above the consensus sequence are a visual aid of conservation (blue, shorter: less conserved; red, higher: more conserved).

Figure S3

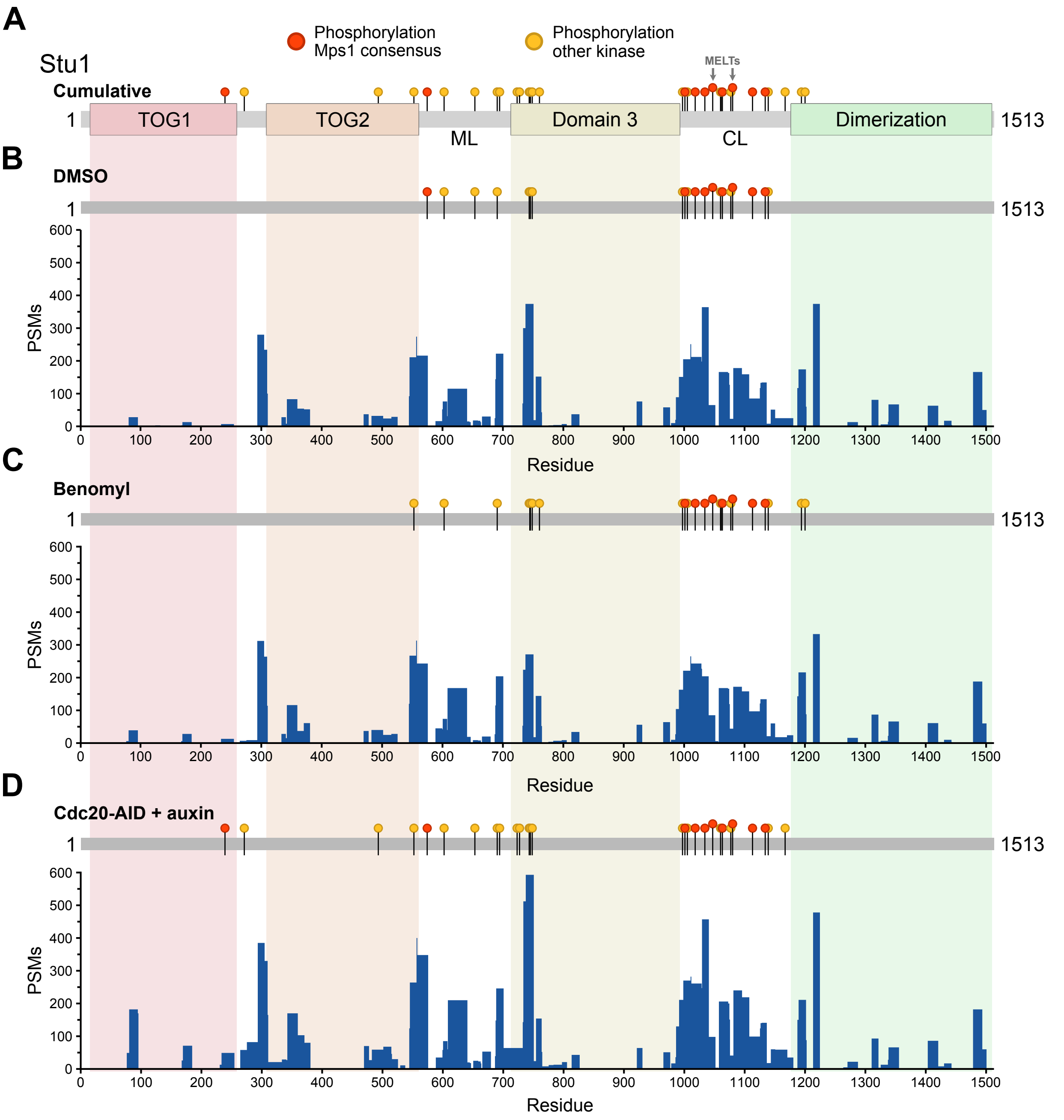

**Figure S3: Mass spectrometry coverage maps of Stu1-3xFLAG immunoprecipitates.** Stu1-3xFLAG immunoprecipitates were analyzed by mass spectrometry for phosphorylation from 3 conditions: asynchronous (DMSO), benomyl-arrested (metaphase arrest, spindle checkpoint active), and *cdc20-AID* arrested (metaphase arrested, spindle checkpoint inactive). The same strain was used for all 3 conditions (*STU1-3xFLAG cdc20-AID leu2::pGPD1-OsTIR1* cells, SBY20662). **(A)** Schematic showing the cumulative phosphorylation detected across the 3 conditions, with colored pins representing each detected phosphorylation site. Red pins indicate sites matching the Mps1 consensus sequence. **(B-D)** Horizontal grey bars represent the positions along the Stu1 protein, with pins representing detected phosphorylation for the indicated sample. Below, a bar graph shows the coverage of each residue per condition, quantified by the number of peptide-spectrum matches (PSMs) that covered each residue for that condition.

Figure S4

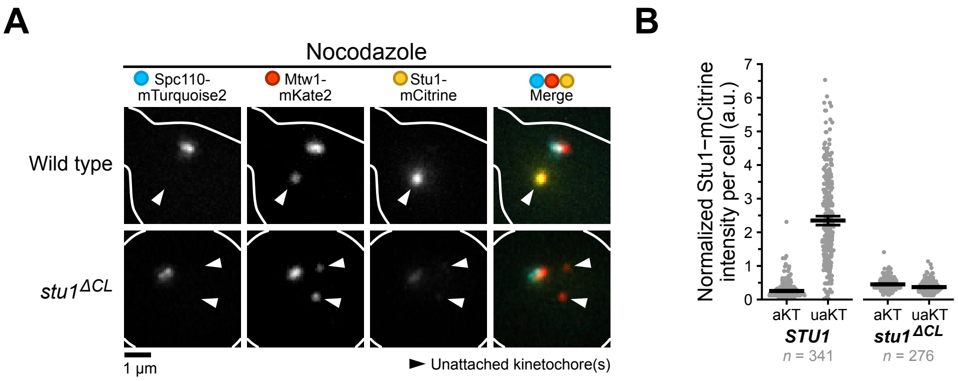

**Figure S4: Deletion of Stu1's CL region disrupts its localization to unattached kinetochores. (A)** *STU1-NLS-mCitrine-3xFLAG* (wild type, SBY22493) and *stu1*$^{\Delta CL}$*-NLS-mCitrine-3xFLAG* (SBY22266) cells containing fluorescent Spc110-mTurquoise2 and Mtw1-mKate2 were released from G1 into 15 µg/mL nocodazole for 1.5 hours to create unattached kinetochores and live imaged. A representative cell is shown for each genotype, and each fluorescent channel is indicated above. **(B)** Quantification of mCitrine fluorescence intensity at attached and unattached kinetochores (aKT and uaKT, respectively) from the experiment shown in (A). mCitrine fluorescence intensities were measured and reported as in Figure 1G. Thick horizontal bars represent the mean value, and error bars represent the 95% confidence interval as determined by bootstrapping, the values of which are shown in **Table S4**. These results are consistent with previous observations[1, 2].

Figure S5

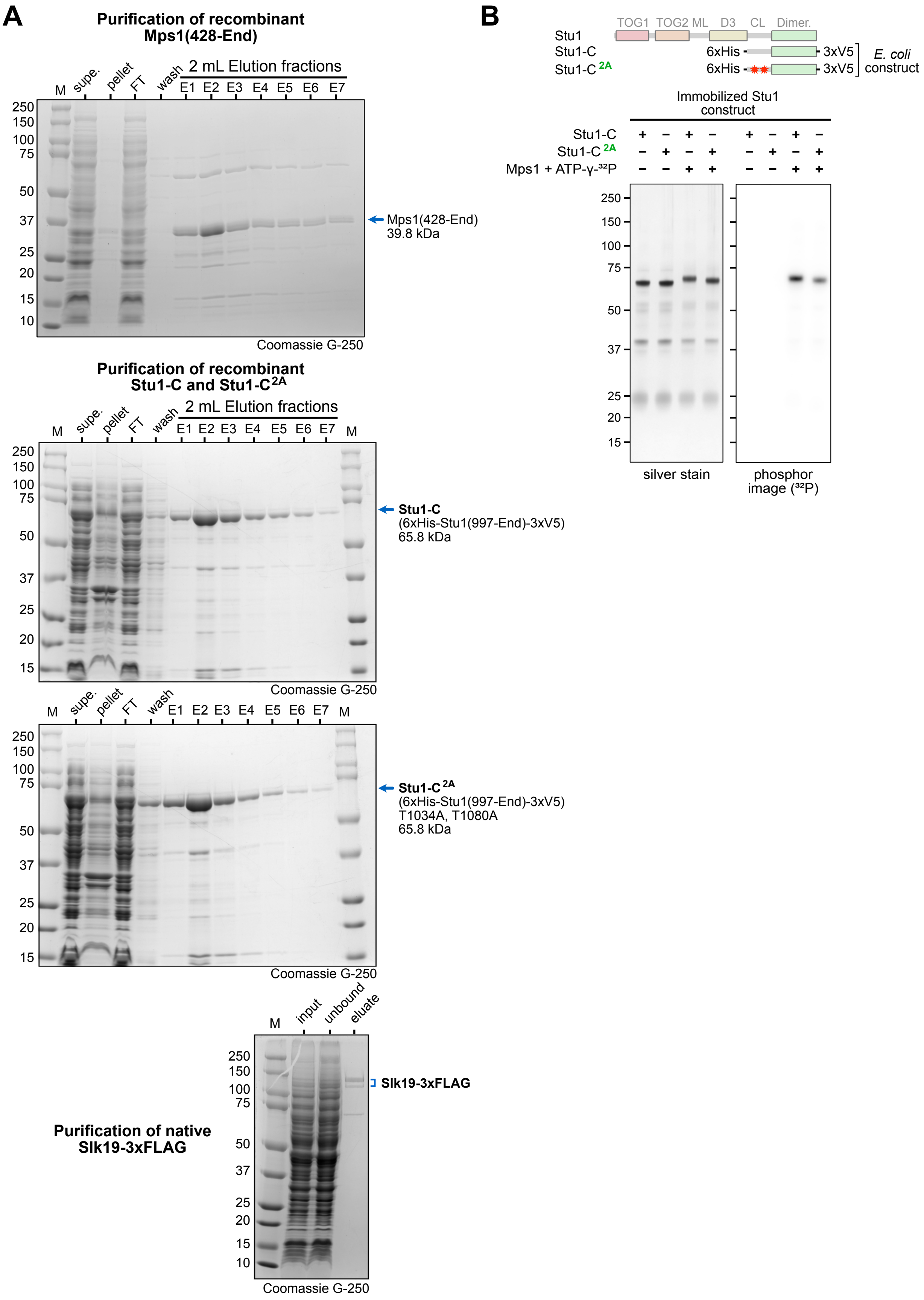
A
Purification of recombinant Mps1(428-End)
M
supe.
pellet
FT
wash
2 mL Elution fractions
E1 E2 E3 E4 E5 E6 E7
250
150
100
75
50
37
25
20
15
10
Mps1(428-End) 39.8 kDa
Coomassie G-250
Purification of recombinant Stu1-C and Stu1-C 2A
Stu1-C (6xHis-Stu1(997-End)-3xV5) 65.8 kDa
Stu1-C 2A (6xHis-Stu1(997-End)-3xV5) T1034A, T1080A 65.8 kDa
Purification of native Slk19-3xFLAG
input
unbound
eluate
Slk19-3xFLAG
B
TOG1 TOG2 ML D3 CL Dimer.
Stu1
Stu1-C
Stu1-C 2A
6xHis
3xV5
E. coli construct
Immobilized Stu1 construct
Mps1 + ATP-γ-32P
silver stain
phosphor image (32P)

**Figure S5: Stu1 is a substrate of the Mps1 kinase *in vitro*. (A)** Coomassie-stained SDS-PAGE gels showing the purification of constructs used in the pulldown experiments. Mps1(428-End), Stu1-C, and Stu1-C$^{2A}$ were expressed and purified from *E. coli*, and native Slk19-3xFLAG was overexpressed and purified from yeast (SBY23482 cells). Abbreviations: M, marker; supe., supernatant; FT, flow-through. **(B)** Mps1 phosphorylates Stu1-C and Stu1-C$^{2A}$ *in vitro*. Top: schematic showing Stu1-C and Stu1-C$^{2A}$ constructs compared to full-length Stu1. Red stars in Stu1-C$^{2A}$ represent phospho-deficient alanine mutations of the MELT motifs. Bottom: recombinant Stu1-C and Stu1-C$^{2A}$ were immobilized on α-V5 beads and incubated with recombinant Mps1 (purified in (A)) with radioactive ATP-$\gamma$-$^{32}$P to directly monitor phosphorylation. SDS-PAGE gels were silver stained (left) and $^{32}$P incorporation was detected on a phosphor screen (right).

#### SUPPLEMENTAL TABLES

**Table S1: Detected phosphorylation sites from Stu1-3xFLAG immunoprecipitation mass spectrometry experiments**

Output of phosphorylation sites with a confidence score $\geq$ 75% and $\geq$ 2 peptide spectrum matches (PSMs) for each phosphorylated peptide across each sample (asynchronous, benomyl-arrested, and metaphase-arrested from *STU1-3xFLAG cdc20-AID leu2::pGPD1-OsTIR1* cells, SBY20662). For each site, the table displays total PSMs, phospho-PSMs, confidences across samples, whether the site has been reported on the Saccharomyces Genome Database (SGD), and the predicted kinase associated with each site.

*[Supplemental Excel Sheet]*

**Table S2: Yeast strains used in this study**

| Strain | Genotype | Figure |
|---|---|---|
| SBY3 | *Mat**a** ura3-1 leu2,3-112 his3-11 trp1-1 ade2-1 can1-100 bar1-1* | Figure 5G, S1C |
| SBY7513 | *Mat**a** ura3-1 leu2,3-112 his3-11 trp1-1 ade2-1 can1-100 bar1-1 slk19Δ::NatMX6* | Figure S1C |
| SBY8956 | *Mat**a** ura3-1 leu2,3-112 his3-11 trp1-1 ade2-1 can1-100 bar1-1 SLK19-13xmyc:HIS3MX6* | Figure 5G |
| SBY8964 | *Mat**a** ura3-1 leu2,3-112 his3-11 trp1-1 ade2-1 can1-100 bar1-1 SLK19-3xFLAG:KanMX6* | Cloning only |
| SBY8968 | *Mat**a** ura3-1 leu2,3-112 his3-11 trp1-1 ade2-1 can1-100 bar1-1 PDS1-18xmyc:LEU2 (pSB205) slk19Δ::NatMX6* | Figure 4D |
| SBY12019 | *Mat**a** ura3-1 leu2,3-112 his3-11 trp1-1 ade2-1 can1-100 bar1-1 STU1-3xV5:KanMX6* | Cloning only |
| SBY12342 | *Mat**a**/Mat**α** ura3-1/ura3-1 leu2,3-112/leu2,3-112 his3-11/his3-11 trp1-1/trp1-1 ade2-1/ade2-1 can1-100/can1-100 bar1-1/bar1-1 DSN1-6xHIS-3xFLAG:URA3/DSN1-6xHIS-3xFLAG:URA3 pSTU2::KanMX:pGAL-Stu2-3V5:HIS3MX6/STU2-13xmyc:HIS3 pGPD1-GAL4(1-848).ER:URA3/pGPD1-GAL4(1-848).ER:URA3* | Cloning only |
| SBY20662 | *Mat**a** ura3-1 leu2::pGPD1-OsTIR1:LEU2 his3-11 trp1-1 ade2-1 can1-100 bar1-1 cdc20-AID:KanMX; Stu1-3FLAG:KanMX6* | Figure 5C, S3 |
| SBY20768 | *Mat**a** ura3-1 leu2,3-112 his3-11 trp1-1 ade2-1 can1-100 bar1-1 STU1-mCitrine:His3MX6* | Cloning only |
| SBY21184 | *Mat**a** ura3-1 leu2,3-112 his3-11 trp1-1 ade2-1 can1-100 bar1-1 MTW1-mKate2:KanMX SPC110-mTurquoise2:TRP1 STU1-mCitrine:HIS3MX6* | Figure 1F, 1G, S1E |
| SBY21197 | *Mat**a** ura3-1 leu2,3-112 his3-11 trp1-1 ade2-1 can1-100 bar1-1 STU1-3xFLAG:hphMX6 BUB1-13xmyc:TRP1* | Figure 1D, S1D |
| SBY21199 | *Mat**a** ura3-1 leu2,3-112 his3-11 trp1-1 ade2-1 can1-100 bar1-1 STU1-3xFLAG:hphMX6* | Figure 1D, S1C |
| SBY21509 | *Mat**a** ura3-1 leu2::pGAL-STU1-mCitrine:LEU2 (pSB3376) his3::pGPD1-OsTIR1 (pSB2273) trp1-1 ade2-1 can1-100 bar1-1 STU1-3HA-IAA7(1-125):KanMX6 MAD1-mTurquoise2:TRP1 MTW1-mKate2:KanMX* | Figure 7B |
| SBY21810 | *Mat**a** ura3-1 leu2,3-112 his3-11 trp1-1 ade2-1 can1-100 bar1-1 stu1-K94S-R95S-3xFLAG:hphMX6 BUB1-13xmyc:TRP1* | Figure 1D, S1D |
| SBY21811 | *Mat**a** ura3-1 leu2,3-112 his3-11 trp1-1 ade2-1 can1-100 bar1-1 stu1-ΔK94-3xFLAG:hphMX6 BUB1-13xmyc:TRP1* | Figure 1D, S1D |
| SBY21935 | *Mat**a** ura3-1 leu2,3-112 his3-11 trp1-1 ade2-1 can1-100 bar1-1 stu1-ΔK94-3xFLAG:hphMX6 SLK19-13xmyc:HIS3MX6* | Figure 1D, 3A |

| | | |
|---|---|---|
| SBY21937 | *Mat**a** ura3-1 leu2,3-112 his3-11 trp1-1 ade2-1 can1-100 bar1-1 stu1-ΔK94-3xFLAG:hphMX6* | Figure S1C |
| SBY21941 | *Mat**a** ura3-1 leu2,3-112 his3-11 trp1-1 ade2-1 can1-100 bar1-1 stu1-K94S-R95S-3xFLAG:hphMX6* | Figure S1C |
| SBY21943 | *Mat**a** ura3-1 leu2,3-112 his3-11 trp1-1 ade2-1 can1-100 bar1-1 stu1-K94S-R95S-3xFLAG:hphMX6 SLK19-13xmyc:HIS3MX6* | Figure 1D |
| SBY21961 | *Mata ura3-1 leu2,3-112 his3-11 trp1-1 ade2-1 can1-100 bar1-1 stu1-ΔK94-3xFLAG:hphMX6 PDS1-18xmyc:LEU2 (pSB205)* | Figure 4D |
| SBY21963 | *Mata ura3-1 leu2,3-112 his3-11 trp1-1 ade2-1 can1-100 bar1-1 MTW1-mKate2:KanMX SPC110-mTurquoise2:TRP1 SLK19-mCitrine:HIS3MX6 stu1-ΔK94-3xFLAG:hphMX6* | Figure 2C, 2D |
| SBY21979 | *Mat**a** ura3-1 leu2,3-112 his3-11 trp1-1 ade2-1 can1-100 bar1-1 STU1-3xFLAG:hphMX6 SLK19-13xmyc:HIS3MX6* | Figure 1D, 3A, 5A, 5B, 5E, 5G |
| SBY21981 | *Mat**a** ura3-1 leu2,3-112 his3-11 trp1-1 ade2-1 can1-100 bar1-1 STU1-3xFLAG:hphMX6 SLK19-13xmyc:HIS3MX6 mps1-1* | Figure 5A |
| SBY22005 | *Mat**a** ura3-1 leu2,3-112 his3-11 trp1-1 ade2-1 can1-100 bar1-1 PDS1-18xmyc:LEU2 (pSB205)* | Figure 4D |
| SBY22043 | *Mat**a** ura3-1 leu2,3-112 his3-11 trp1-1 ade2-1 can1-100 bar1-1 MTW1-mKate2:KanMX SPC110-mTurquoise2:TRP1 SLK19-mCitrine:HIS3MX6 STU1-3xFLAG:hphMX6* | Figure 2C, 2D, 6A, 6B |
| SBY22050 | *Mat**a** ura3-1 leu2::pSTU1-stu1ΔTOG1(Δ2-260)-NLS-mCitrine-3xFLAG:LEU2 (pSB3390) his3-11 trp1-1 ade2-1 can1-100 bar1-1 stu1Δ::NatMX6 SLK19-13xmyc:HIS3MX6 BUB1-3xV5:KAN* | Figure 1A |
| SBY22056 | *Mat**a** ura3-1 leu2::pSTU1-stu1ΔCL(Δ997-1180)-NLS-mCitrine-3xFLAG:LEU2 (pSB3417) his3-11 trp1-1 ade2-1 can1-100 bar1-1 stu1Δ::NatMX6 SLK19-13xmyc:HIS3MX6 BUB1-3xV5:KAN* | Figure 1A, 5F |
| SBY22069 | *Mat**a** ura3-1 leu2::pSTU1-STU1-NLS-mCitrine-3xFLAG:LEU2 (pSB3379) his3-11 trp1-1 ade2-1 can1-100 bar1-1 stu1Δ::NatMX6 SLK19-13xmyc:HIS3MX6 BUB1-3xV5:KAN* | Figure 1A, 5F |
| SBY22266 | *Mat**a** ura3-1 leu2::pSTU1-stu1ΔCL(Δ997-1180)-NLS-mCitrine-3xFLAG:LEU2 (pSB3417) his3-11 trp1-1 ade2-1 can1-100 bar1-1 stu1Δ::NatMX6 MTW1-mKate2:KanMX SPC110-mTurquoise2:TRP1* | Figure S4 |
| SBY22330 | *Mat**a** ura3-1 leu2,3-112 his3-11 trp1-1 ade2-1 can1-100 bar1-1 STU1-3xFLAG:hphMX6 SLK19-13xmyc:HIS3MX6 cdc5-1* | Figure 5A |
| SBY22334 | *Mat**a** ura3-1 leu2,3-112 his3-11 trp1-1 ade2-1 can1-100 bar1-1 PDS1-18xmyc:LEU2 (pSB205) mad3Δ::KanMX6* | Figure 4D |
| SBY22476 | *Mat**a** ura3-1 leu2,3-112 his3-11 trp1-1 ade2-1 can1-100 bar1-1 stu1-T1080A-3xFLAG:hphMX6 SLK19-13xmyc:HIS3MX6* | Figure 5E, 5G |

| | | |
|---|---|---|
| SBY22493 | *Mat**a** ura3-1 leu2::pSTU1-STU1-NLS-mCitrine-3xFLAG:LEU2 (pSB3379) his3-11 trp1-1 ade2-1 can1-100 bar1-1 MTW1-mKate2:KanMX SPC110-mTurquoise2:TRP1 stu1Δ::NatMX6* | Figure 6C, 6D, S4 |
| SBY22568 | *Mat**a** ura3-1 leu2,3-112 his3-11 trp1-1 ade2-1 can1-100 bar1-1 MTW1-mKate2:KanMX SPC110-mTurquoise2:TRP1 stu1-ΔK94-mCitrine:HIS3MX6* | Figure S1E |
| SBY23112 | *Mat**a** ura3-1 leu2::pSTU1-stu1-T1034A-T1080A-NLS-mCitrine-3xFLAG:LEU2 (pSB3518) his3-11 trp1-1 ade2-1 can1-100 bar1-1 BUB1-3xV5:KAN SLK19-13xMyc:HIS stu1Δ::NatMX6* | Figure 5F |
| SBY23135 | *Mat**a** ura3-1 leu2::pSTU1-stu1-T1034A-T1080A-NLS-mCitrine-3xFLAG:LEU2 (pSB3518) his3-11 trp1-1 ade2-1 can1-100 bar1-1 MTW1-mKate2:KanMX SPC110-mTurquoise2:TRP1 stu1Δ::NatMX6* | Figure 6C, 6D |
| SBY23370 | *Mat**a** ura3-1 leu2,3-112 his3-11 trp1-1 ade2-1 can1-100 bar1-1 MTW1-3xmYPet:CaURA3 (pSB3562) SPC110-mTurquoise2:TRP1* | Figure 4B |
| SBY23372 | *Mat**a** ura3-1 leu2,3-112 his3-11 trp1-1 ade2-1 can1-100 bar1-1 MTW1-3xmYPet:CaURA3 (pSB3562) SPC110-mTurquoise2:TRP1 slk19Δ::NatMX6* | Figure 2A, 2B |
| SBY23478 | *Mat**a** ura3-1 leu2,3-112 his3-11 trp1-1 ade2-1 can1-100 bar1-1 MTW1-3xmYPet:CaURA3 (pSB3562) SPC110-mTurquoise2:TRP1 mps1-1* | Figure 4B |
| SBY23480 | *Mat**a** ura3-1 leu2,3-112 his3-11 trp1-1 ade2-1 can1-100 bar1-1 MTW1-3xmYPet:CaURA3 (pSB3562) SPC110-mTurquoise2:TRP1 STU1-3xFLAG:hphMX6* | Figure 2A, 2B, 3B, 6E, 6F |
| SBY23482 | *Mat**a** ura3-1 leu2::pGAL-SLK19-3xFLAG:LEU2 (pSB3515) his3-11 trp1-1 ade2-1 can1-100 bar1-1 SLK19-3xFLAG:KanMX6 mps1-1* | Figure S4 |
| SBY23484 | *Mat**a** ura3-1 leu2,3-112 his3-11 trp1-1 ade2-1 can1-100 bar1-1 MTW1-3xmYPet:CaURA3 (pSB3562) SPC110-mTurquoise2:TRP1 stu1-ΔK94-3xFLAG:hphMX6* | Figure 2A, 2B |
| SBY23547 | *Mata ura3-1 leu2,3-112 his3::pGPD1-OsTIR1 (pSB2273) trp1-1 ade2-1 can1-100 bar1-1 MTW1-3xmYPet:CaURA3 (pSB3562) SPC110-mTurquoise2:TRP1 cdc20-AID:NatMX6 BUB3-3xV5-IAA7(1-125):hphMX6* | Figure 4C |
| SBY23751 | *Mat**a** ura3-1 leu2,3-112 his3-11 trp1-1 ade2-1 can1-100 bar1-1 stu1-T1034A-T1080A-3xFLAG:hphMX6 SLK19-13xmyc:HIS3MX6* | Figure 5E, 5G |
| SBY23753 | *Mat**a** ura3-1 leu2,3-112 his3-11 trp1-1 ade2-1 can1-100 bar1-1 MTW1-3xmYPet:CaURA3 (pSB3562) SPC110-mTurquoise2:TRP1 stu1-T1034A-T1080A-3xFLAG:hphMX6* | Figure 6E, 6F |
| SBY23770 | *Mat**a** ura3-1 leu2,3-112 his3-11 trp1-1 ade2-1 can1-100 bar1-1 stu1-T1034A-3xFLAG:hphMX6 SLK19-13xmyc:HIS3MX6* | Figure 5E, 5G |

| | | |
|---|---|---|
| SBY23772 | *Mat**a** ura3-1 leu2,3-112 his3-11 trp1-1 ade2-1 can1-100 bar1-1 MTW1-3xmYPet:CaURA3 (pSB3562) SPC110-mTurquoise2:TRP1 stu1-T1034A-3xFLAG:hphMX6* | Figure 6E, 6F |
| SBY23822 | *Mat**a** ura3-1 leu2,3-112 his3-11 trp1-1 ade2-1 can1-100 bar1-1 MTW1-3xmYPet:CaURA3 (pSB3562) SPC110-mTurquoise2:TRP1 stu1-T1080A-3xFLAG:hphMX6* | Figure 6E, 6F |
| SBY23897 | *Mat**a** ura3-1 leu2,3-112 his3-11 trp1-1 ade2-1 can1-100 bar1-1 STU1::BUB3-STU1-3xFLAG:hphMX6 SLK19-13xmyc:HIS3MX6* | Figure 3A |
| SBY23901 | *Mat**a** ura3-1 leu2,3-112 his3-11 trp1-1 ade2-1 can1-100 bar1-1 STU1::BUB3-stu1-ΔK94-3xFLAG:hphMX6 SLK19-13xmyc:HIS3MX6* | Figure 3A |
| SBY24036 | *Mat**a** ura3-1 leu2,3-112 his3-11 trp1-1 ade2-1 can1-100 bar1-1 MTW1-3xmYPet:CaURA3 (pSB3562) SPC110-mTurquoise2:TRP1 STU1::BUB3-STU1-3FLAG:hphMX6* | Figure 3B |
| SBY24038 | *Mat**a** ura3-1 leu2,3-112 his3-11 trp1-1 ade2-1 can1-100 bar1-1 MTW1-3xmYPet:CaURA3 (pSB3562) SPC110-mTurquoise2:TRP1 STU1::BUB3-stu1-ΔK94-3FLAG:hphMX6* | Figure 3B |
| SBY24246 | *Mat**a** ura3-1 leu2,3-112 his3-11 trp1-1 ade2-1 can1-100 bar1-1 MTW1-mKate2:KanMX SPC110-mTurquoise2:TRP1 SLK19-mCitrine:HIS3MX6 stu1-T1034A-T1080A-3xFLAG:hphMX6* | Figure 6A, 6B |
| SBY24326 | *Mat**a** ura3-1 leu2,3-112 his3-11 trp1-1 ade2-1 can1-100 bar1-1 MTW1-mKate2:KanMX SPC110-mTurquoise2:TRP1 SLK19-mCitrine:HIS3MX6 stu1-T1034A-3xFLAG:hphMX6* | Figure 6A, 6B |
| SBY24331 | *Mat**a** ura3-1 leu2,3-112 his3-11 trp1-1 ade2-1 can1-100 bar1-1 MTW1-mKate2:KanMX SPC110-mTurquoise2:TRP1 SLK19-mCitrine:HIS3MX6 stu1-T1080A-3xFLAG:hphMX6* | Figure 6A, 6B |
| SBY24733 | *Mat**a** ura3-1 leu2,3-112 his3-11 trp1-1 ade2-1 can1-100 bar1-1 MTW1-mKate2:KanMX SPC110-mTurquoise2:TRP1 stu1-ΔK94-mCitrine:HIS3MX6* | Figure 1F, 1G |

All yeast strains are derivatives of the W303 background (SBY3). Stably integrated plasmids are indicated in brackets.

**Table S3: Plasmids used in this study.**

| Plasmid | Description |
|---|---|
| pSB54 | *KanMX6* (Gene deletions) |
| pSB63 | *13xmyc*, *TRP1* (C-terminal tagging) |
| pSB64 | *13xmyc*, *HIS3MX6* (C-terminal tagging) |
| pSB205 | *PDS1-18xMyc*, *LEU2* (*PDS1* integrating, HindIII) (A gift from Kim Nasmyth) |
| pSB637 | *NatMX6* (Gene deletions) |
| pSB813 | *3xFLAG*, *hphMX6* (C-terminal tagging) |
| pSB2046 | *3xV5*, *KanMX* (C-terminal tagging, a gift from Angelika Amon, p1726) |
| pSB2068 | pFA6a-*3xV5*, *KanMX6* (C-terminal tagging, a gift from Karsten Weis, pL264) |
| pSB2223 | pNH605(XbaI)-pDMC1, *LEU2* (pL300, a gift from Karsten Weis) |
| pSB2273 | *pGPD1-OsTIR1*, *HIS3* (*HIS3* integrating, PmeI) (A gift from Karsten Weis, pL253) |
| pSB2933 | pFA6a-*mKate2*, *KanMX* (C-terminal tagging, a gift from Elçin Ünal) |
| pSB2936 | pFA6a-*mCitrine*, *HIS3MX6* (C-terminal tagging, a gift from Elçin Ünal) |
| pSB2940 | pFA6a-*mTurquoise2*, *TRP1* (C-terminal tagging, a gift from Elçin Ünal) |
| pSB3218 | *pPGK1-SpCas9,* GFP-sgRNA parent vector, *URA3* (A gift from Elçin Ünal) |
| pSB3363 | *pSTU1-STU1-3xFLAG*, *LEU2* (*LEU2* integrating, PmeI) |
| pSB3376 | *pGAL-STU1-mCitrine*, *LEU2* (*LEU2* integrating, PmeI) |
| pSB3379 | *pSTU1-STU1-NLS-mCitrine-3xFLAG*, *LEU2* (*LEU2* integrating, PmeI) |
| pSB3390 | *pSTU1-STU1-ΔTOG1(Δ2-260)-NLS-mCitrine-3xFLAG*, *LEU2* (*LEU2* integrating, PmeI) |
| pSB3409 | *3xV5-IAA7(1-125)*, *hphMX6* (C-terminal tagging) |
| pSB3417 | *pSTU1-STU1-ΔCL(Δ997-1180)-NLS-mCitrine-3xFLAG*, *LEU2* (*LEU2* integrating, PmeI) |
| pSB3419 | *pPGK1-SpCas9*, sgRNA targeting Stu1 TOG1 basic patch (TTGTTACCTTATCAAACGTG), *URA3* |
| pSB3465 | *pPGK1-SpCas9*, sgRNA targeting Stu1 N-terminal region for *BUB3* insertion (ATTATTGGTCTCATTGTTGA), *URA3* |
| pSB3483 | *pPGK1-SpCas9*, sgRNA targeting Stu1 T1034 and T1080 (CCTCCTTTGCTGGGTTTGGC), *URA3* |
| pSB3515 | *pGAL-SLK19-3xFLAG*, *LEU2* (*LEU2* integrating, PmeI) |
| pSB3518 | *pSTU1-stu1-T1034A-T1080A-NLS-mCitrine-3xFLAG*, *LEU2* (*LEU2* integrating, PmeI) |
| pSB3519 | pET-21b-*STU1-3xV5-6xHis* (cloning for recombinant expression) |
| pSB3524 | pET-21b-*STU1(CL+D4)-3xV5-6xHis* (cloning for recombinant expression) |
| pSB3554 | pFA6a-link-*ymYPET*, *URA3* (Addgene, #168056) (for cloning 3xmYPet) |
| pSB3559 | pET-21b-*STU1(CL+D4)-3xV5* (cloning for recombinant expression) |

| | |
|---|---|
| pSB3560 | 6xHis-Stu1-C-3xV5 (recombinant expression of Stu1-C, a pET-21b(+) derivative) |
| pSB3561 | pFA6a-link-*3xymYPET*, *URA3* (precursor to pSB3562) |
| pSB3562 | *MTW1-3xmYPet*, *URA3* (C. albicans) (*MTW1* integrating, SnaBI) |
| pSB3564 | 6xHis-Stu1-C-2A-3xV5 (recombinant expression of Stu1-C-2A, a pET-21b(+) derivative) |
| pSB3587 | Lambda PPase (pCDFDuet1, a gift from Arshad Desai) |
| pSB3593 | 6xHis-Mps1(full-length) Lambda PPase (pCDFDuet1 derivative) |
| pSB3601 | 6xHis-Mps1(428-End) Lambda PPase (pCDFDuet1 derivative) |

Integration plasmids with the restriction enzyme used for linearization are noted in parentheses.

**Table S4: Statistics of image quantification**

| | Sample | Location | Mean | 95% CI (lower bound) | 95% CI (upper bound) |
|---|---|---|---|---|---|
| **Figure 1G** | *STU1* | aKT | 0.20 | 0.17 | 0.23 |
| | *STU1* | uaKT | 2.31 | 2.11 | 2.51 |
| | *stu1-ΔK94* | aKT | 0.33 | 0.32 | 0.35 |
| | *stu1-ΔK94* | uaKT | 0.34 | 0.32 | 0.36 |
| | **Sample** | **Mean** | **p-value** | | |
| **Figure 2B** | *Wild type* | 2.61 | 2.2E-16 | | |
| | *slk19Δ* | 4.83 | | | |
| | *STU1-3xFLAG* | 2.68 | 2.2E-16 | | |
| | *stu1-ΔK94-3xFLAG* | 3.98 | | | |
| | **Sample** | **Location** | **Mean** | **95% CI (lower bound)** | **95% CI (upper bound)** |
| **Figure 2D** | *STU1* | aKT | 0.40 | 0.37 | 0.43 |
| | *STU1* | uaKT | 4.75 | 4.45 | 5.08 |
| | *stu1-ΔK94* | aKT | 0.67 | 0.64 | 0.70 |
| | *stu1-ΔK94* | uaKT | 1.12 | 0.98 | 1.30 |
| | **Sample** | **Mean** | **p-value** | | |
| **Figure 3B** | *STU1* | 2.65 | 0.835 | | |
| | *BUB3-STU1* | 2.67 | | | |
| | *STU1* | 2.65 | 0.085 | | |
| | *BUB3-stu1-ΔK94* | 2.77 | | | |
| | **Sample** | **Mean** | **p-value** | | |
| **Figure 3B** | *Wild type* | 3.33 | 2.2E-16 | | |
| | *mps1-1* | 5.17 | | | |
| | **Condition** | **Mean** | **p-value** | | |
| **Figure 3C** | DMSO | 2.52 | 0.692 | | |
| | Auxin | 2.47 | | | |

| | | | | | |
|---|---|---|---|---|---|
| **Figure 6B** | **Sample** | **Location** | **Mean** | **95% CI (lower bound)** | **95% CI (upper bound)** |
| | *STU1* | aKT | 0.42 | 0.38 | 0.46 |
| | *STU1* | uaKT | 5.27 | 4.90 | 5.67 |
| | *stu1-T1034A* | aKT | 0.67 | 0.63 | 0.71 |
| | *stu1-T1034A* | uaKT | 2.40 | 2.17 | 2.67 |
| | *stu1-T1080A* | aKT | 0.57 | 0.52 | 0.62 |
| | *stu1-T1080A* | uaKT | 4.22 | 3.93 | 4.53 |
| | *stu1-2A* | aKT | 0.78 | 0.75 | 0.81 |
| | *stu1-2A* | uaKT | 0.89 | 0.84 | 0.94 |
| **Figure 6D** | **Sample** | **Location** | **Mean** | **95% CI (lower bound)** | **95% CI (upper bound)** |
| | *STU1* | aKT | 0.22 | 0.18 | 0.26 |
| | *STU1* | uaKT | 2.74 | 2.56 | 2.92 |
| | *stu1-2A* | aKT | 0.34 | 0.32 | 0.37 |
| | *stu1-2A* | uaKT | 0.42 | 0.38 | 0.45 |
| **Figure 6F** | **Sample** | **Mean** | **p-value** | | |
| | *STU1-3xFLAG* | 2.72 | 0.071 | | |
| | *stu1-T1034A-3xFLAG* | 2.84 | | | |
| | *STU1-3xFLAG* | 2.72 | 0.1812 | | |
| | *stu1-T1080A-3xFLAG* | 2.58 | | | |
| | *STU1-3xFLAG* | 2.72 | 2.2E-16 | | |
| | *stu1-2A-3xFLAG* | 3.86 | | | |
| **Figure S4** | **Sample** | **Location** | **Mean** | **95% CI (lower bound)** | **95% CI (upper bound)** |
| | *STU1* | aKT | 0.25 | 0.23 | 0.28 |
| | *STU1* | uaKT | 2.35 | 2.22 | 2.48 |
| | *stu1-ΔCL* | aKT | 0.45 | 0.43 | 0.47 |
| | *stu1-ΔCL* | uaKT | 0.37 | 0.35 | 0.39 |

Output of statistical analyses including means, p-values, and 95% confidence intervals arranged by figure.

**Table S5: Regular expressions used to predict kinases in phosphorylation data**

<table>
<tr><th>Kinase</th><th>Regular expression used</th><th>References</th></tr>
<tr><td>Mps1</td><td>[DENCQ][A-Z]([ST])[^PN]</td><td>3, 4</td></tr>
<tr><td>Mps1 (if -2 phos)</td><td>[ST][A-Z]([ST])[^PN]</td><td>5</td></tr>
<tr><td>Ipl1</td><td>[RK][A-Z]([ST])[^P]</td><td>3, 6</td></tr>
<tr><td>Cdk1</td><td>([ST])[P](?:[A-Z][KR])?</td><td>7</td></tr>
<tr><td>Cdc5</td><td>[DENCQ][A-Z]([ST])(?:[^PDEN]|[FMYI])</td><td>3, 8, 9</td></tr>
<tr><td>Cdc5 (if +1 phos)</td><td>[DENCQ][A-Z]([ST])[ST]</td><td>3</td></tr>
</table>

These regular expressions were used to match common mitotic kinase consensus motifs to a given phosphorylation site in **Table S1**. The corresponding references describing each consensus motif are listed.

**Table S6: Oligonucleotides and gBlocks used in this study**

| Sequence ID | Purpose | Sequence (5' to 3') |
|---|---|---|
| SB53 | Forward genotyping primer to check C-terminal tags containing TEF terminator (*SLK19* deletion, *STU1* deletion) | TTCGCCTCGACATCATCTGC |
| SB249 | Forward primer for *SLK19* C-terminal tagging with *13xmyc:HIS3MX6* and *mCitrine:HIS3MX6* | GAGAAGAAAAGCAGGAGTTACTCAAGTTGTTAGAAAATGAAAAAAAAGGTCGACGGATCCCCGGGTT |
| SB250 | Reverse primer for *SLK19* C-terminal tagging with *13xmyc:HIS3MX6* and *mCitrine:HIS3MX6* | CTCATGACATATTAAGGGAAAAGATAAAATGCAAAAGAAAAAAATGCGTTCGATGAATTCGAGCTCGTT |
| SB251 | Reverse genotyping primer for *SLK19* C-terminal tagging and deletions | GCACACTGAAATCTAATAGTTG |
| SB392 | Reverse primer for *MTW1* C-terminal tagging (mKate2:KanMX) | ATACATCATATCATAGCACATACTTTTTCCCACTTTATATCGATGAATTCGAGCTCGTT |
| SB549 | Forward primer for *MAD3* gene replacement with *KanMX6* (deletion) | GTAAACAAAATCATGCGAAAATACAATAAAAGACGTTAACTTGATAGAGGTCGACGGATCCCCGGGTT |
| SB550 | Reverse primer for *MAD3* gene replacement with *KanMX6* (deletion) | GTTTACGATTGGCCAGTATACTTACTCATTCATGGGATTAGTT |
| SB552 | Reverse primer for *BUB1* C-terminal tagging (*3xV5-IAA7(1-125):hphMX6*) | GGCAGGACACCAAAAAGTCACCTATGCGGGAGATGAAGGCATATTTATTCATCGATGAATTCGAGCTCGTT |
| SB557 | Forward genotyping primer to check *MAD3* deletion | GCTAAATGAATACACAGCGTG |
| SB558 | Reverse genotyping primer to check *MAD3* deletion | CGCTTAATGCAGGGAGCTC |
| SB800 | Forward primer for *BUB1* C-terminal tagging (*3xV5-IAA7(1-125):hphMX6*) | GAGGAGTTATCACATTTTCAATATAAGGGGAAACCGTCAAGGAGATTTGGTCGACGGATCCCCGGGTT |
| SB807 | Reverse genotyping primer for *BUB1* C-terminal tagging | CATAAGTGACAGATGTCAAG |

| | | |
|---|---|---|
| SB1222 | Forward primer for *SLK19* gene replacement with *NatMX6* (deletion) | CACCCAGTTAAAAAAGGTTTTGAGCACATATCGTAATTCCGGATCCCCGGGTTAATTAA |
| SB1223 | Reverse primer for *SLK19* gene replacement with *NatMX6* (deletion) | TATTAAGGGAAAAGATAAAATGCAAAAGAAAAAAATGCGTTCGATGAATTCGAGCTCGTT |
| SB2721 | Forward primer for *STU1* C-terminal tagging with *mCitrine:HIS3MX6* | TTATTCGATTGTTTGCCTAAGAATGTCTTTAAAATGATCATGTTCATCGCCTCAAACGAAGGTCGACGGATCCCCGGGTT |
| SB2722 | Reverse primer for *STU1* gene replacement with NatMX6 (deletion) and *STU1* C-terminal tagging with *mCitrine:HIS3MX6* | AAGAAACTCTGGTGAGACGCGTCACGGTAAAAAAAAATTACGCGTCTACCACGCTATTCTTCGATGAATTCGAGCTCGTT |
| SB2725 | Forward primer for *SLK19* C-terminal tagging with *3xFLAG:HphMX6* | CTGTCTTCAGAAAGAGAAGAAAAGCAGGAGTTACTCAAGTTGTTAGAAAATGAAAAAAAAAGGGAACAAAAGCTGGAGCT |
| SB2726 | Reverse primer for *SLK19* C-terminal tagging with *3xFLAG:HphMX6* | AATATTTTATTCTCATGACATATTAAGGGAAAAGATAAAATGCAAAAGAAAAAAATGCGTCTATAGGGCGAATTGGGT |
| SB2727 | Forward genotyping primer for *SLK19* C-terminal tagging | GCAGAAGATTTGTATATCCAG |
| SB2728 | Reverse genotyping primer for *SLK19* C-terminal tagging | CATGGACCGCAATGTCTTTG |
| SB2729 | Forward primer for *STU1* C-terminal tagging with *3xFLAG:HphMX6* | TTATTCGATTGTTTGCCTAAGAATGTCTTTAAAATGATCATGTTCATCGCCTCAAACGAAAGGGAACAAAAGCTGGAGCT |
| SB2730 | Reverse primer for *STU1* C-terminal tagging with *3xFLAG:HphMX6* | AAGAAACTCTGGTGAGACGCGTCACGGTAAAAAAAAATTACGCGTCTACCACGCTATTCTCTATAGGGCGAATTGGGT |
| SB2967 | Forward primer to amplify *STU1-3xV5* from gDNA for cloning. | CGTTACCCCGTCGGCGC |
| SB3118 | Reverse primer to sequence *stu1 TOG1* CRISPR mutagenesis and *BUB3-STU1* CRISPR insertion. | GGCTTTAATGGAAGCCAGCC |
| SB3255 | Forward genotyping primer for *BUB1* C-terminal tagging | GGAAAAAAGAAATATGGGGCG |

| | | |
|---|---|---|
| SB3794 | Forward genotyping primer to check *STU1* C-terminal tagging | GGTTACTGAAATTCAGCC |
| SB3795 | Reverse genotyping primer to check *STU1* C-terminal tagging and *STU1* deletions | GGCTTGTCTGCATCAGGC |
| SB4156 | Forward primer to C-terminally tag *BUB1* with *3xV5:KAN* | CTAAGCATTGAAGAGGAGTTATCAC ATTTTCAATATAAGGGGAAACCGTCA AGGAGATTTGCGGCCGCTCTAGAA CTAGTGG |
| SB4157 | Reverse primer to C-terminally tag *BUB1* with *3xV5:KAN* | ATGGAATCTGGCAGGACACCAAAAA GTCACCTATGCGGGAGATGAAGGC ATATTTATTCACCCCCTCGAGGTCG ACGGTATCG |
| SB4373 | Forward primer to amplify *pGAL-STU2* | GATCGATCGGGCCCGTAAAGAGCC CCATTATCTTAG |
| SB4374 | Reverse primer to amplify *pGAL-STU2* | GATCGATCCTCGAGTTATGGATCTGT ACTATCCAGTCC |
| SB6527 | Forward genotyping primer for *SPC110* C-terminal tags | TCACTTCCAGACGATGATGAACT |
| SB6528 | Reverse genotyping primer for *SPC110* C-terminal tags | ACCACATACATAGATATACCCTACG T |
| SB6842 | Forward primer for *MTW1* C-terminal tagging (*mKate2:KanMX*) | TATTGAAGAGCCTCAATTGGATTTAC TTGATGATGTGTTAGGTCGACGGATC CCCGGGTT |
| SB7213 | Forward primer for *SPC110* C-terminal tagging (mTurquoise2:TRP1) | GATAACAGATTGCGAATACTAAGAG ATAGAATTGAGAGTAGCAGCGGGC GTATATCTTGGCGGATCCCCGGGTT AATTAA |
| SB7214 | Reverse primer for SPC110 C-terminal tagging (*mTurquoise2:TRP1*) | ATGATAGAGTAAGCGATAGAATTCGA GCTCGTTTAAAC |
| SB7555 | Forward primer to sequence *BUB3-STU1* CRISPR insertion. Forward primer to amplify *pSTU1-STU1-3xFLAG* | TGAACTCTGCGTTGTCATCAACATC G |
| SB7557 | Reverse primer to amplify *pSTU1-STU1-3xFLAG* | AATTAACCCGGGGATCCGTCGACC |
| SB7558 | To amplify *LEU2* integration vector backbone with homology to *pSTU1* for Gibson assembly | GCGTCGATGTTGATGACAACGCAGA GTTCAGGGCCCGGTACCGTTCGTTT CC |

| | | |
|---|---|---|
| SB7559 | To amplify integration vector backbone with homology to *pSTU1* for Gibson assembly | GCTGCAGGTCGACGGATCCCCGGGTTAATTCCACCGCGGTGGAGCTCTAAGC |
| SB7601 | To linearize backbone of *pSTU1-STU1-3FLAG* integration vector in between *STU1* and *3xFLAG* | GAAAGGGAACAAAAGCTGGAGCTCG |
| SB7602 | To linearize backbone of *pSTU1-STU1-3FLAG* integration vector in between *STU1* and *3xFLAG*. Also to amplify C-terminus of Stu1 to generate stu1-2A in pSB3564. | TTCGTTTGAGGCGATGAACATGATC |
| SB7603 | To amplify *LEU2* integration vector backbone containing the pGAL1-10 promoter. Contains homology to STU1-mCitrine. | ACCCATGGTATTGATGAATTGTACAAATAACTCGAGGGAGCAAGGCAGG |
| SB7604 | To amplify LEU2 integration vector backbone containing the pGAL1-10 promoter. Contains homology to *STU1-mCitrine*. | ATTATTGGTCTCATTGTTGAAGGACGACATTTTGAGATCCGGGTTTTTTCTCC |
| SB7605 | Forward primer to sequence s*tu1 TOG1* CRISPR mutagenesis | ATGTCGTCCTTCAACAATGAGACC |
| SB7605 | To amplify *STU1-mCitrine* from gDNA for Gibson assembly. | ATGTCGTCCTTCAACAATGAGACC |
| SB7606 | To amplify *STU1-mCitrine* from gDNA for Gibson assembly. | TTATTTGTACAATTCATCAATACCATGGG |
| SB7611 | Forward to amplify SV40 NLS + mCitrine to generate *pSTU1-STU1-NLS-mCitrine-3xFLAG* | AAAATGATCATGTTCATCGCCTCAAACGAACCAAAGAAGAAGAGAAAGGTTGGTCGACGGATCCCCGGGTT |
| SB7612 | Reverse to amplify SV40 NLS + mCitrine to generate *pSTU1-STU1-NLS-mCitrine-3xFLAG* | ATAATCGAGCTCCAGCTTTTGTTCCCTTTCTTTGTACAATTCATCAATACCATGGG |
| SB7642 | Forward primer to delete TOG1 from *pSTU1-STU1-NLS-mCitrine-3FLAG* | AGTTTAGCAAAGTCACAGGACC |
| SB7643 | Reverse primer to delete TOG1 from pSTU1-STU1-NLS-mCitrine-3FLAG | CATTATTTCTGAAGAATACAAGGTCTCTCC |
| SB7674 | Reverse primer to amplify *STU1-3xV5* from gDNA for cloning. | GACTGTCAAGGAGGGTATTCTGG |

| | | |
|---|---|---|
| SB7831 | Reverse primer to delete CL from *pSTU1-STU1-NLS-mCitrine* | AATGATGTCTTCTACCTTAAAGTTGGC |
| SB7832 | Forward primer to delete CL from *pSTU1-STU1-NLS-mCitrine* | CAAAGGATCACAGTAAAGAGAGAAAGAG |
| SB7852 | Protospacer (forward) with overhang for ligation into BsmBI-linearized pSB3218 to generate pSB3419 for Stu1 TOG1 CRISPR mutagenesis | GACTTTGTTACCTTATCAAACGTG |
| SB7853 | Protospacer (reverse) with overhang for ligation into BsmBI-linearized pSB3218 to generate pSB3419 for Stu1 TOG1 CRISPR mutagenesis | AAACCACGTTTGATAAGGTAACAA |
| SB7855 | CRISPR homology-directed repair template to generate *stu1-K94S-R95S* | TTATTTCACTGCGTTACTGTTCATCTCCGGCCATTACGCTTACCGTTCGTACCCGCGGTTAATCTTCCTATCACATTCCTCGCTTTGTTACCTTATCTCTTCTGTGGCCATGCAGTCTCCAGTACAATTCAATGACACTCTAGTTGAGCAATTACTAAACCACTTAATTTTCGAGTTGCCTAATGAGAAGAAATTTTGGC |
| SB7875 | Forward primer for *STU1* gene replacement with *NatMX6* (deletion) | CAGGCATATTTAGCGGTAATTATTAGGGTTTTTGGAGAGACCTTGTATTCTTCAGAAATACGGATCCCCGGGTTAATTAA |
| SB7961 | To generate 6xHis tag at Stu1 N-terminus. | ATGTATATCTCCTTCTTAAAGTTAAACAAAATTATTTC |
| SB7990 | Protospacer (forward) with overhang for ligation into BsmBI-linearized pSB3218 to generate pSB3483 for Stu1 T1034 and T1080 CRISPR editing | GACTCCTCCTTTGCTGGGTTTGGC |
| SB7991 | Protospacer (reverse) with overhang for ligation into BsmBI-linearized pSB3218 to generate pSB3483 for Stu1 T1034 and T1080 CRISPR editing | AAACGCCAAACCCAGCAAAGGAGG |
| SB7993 | CRISPR homology-directed repair template to generate *stu1-T1080A*. | ATCAAGCACAGACAGCGTAGTTATTCATGATGATAATGACAAAGATAAAAAGCTTTCAGAAATGGCTAAAATAGTAAGTGTTTATCAACTGGATCAGCCAAA |

| | | |
|---|---|---|
| | | CCCAGCAAAGGAGGAAGATGATATAGATATGGAAAATTCTCAAAAATCTGATTTGAATTTAAGTGAAATTTTTCAAAACAGTGGTGAAAATACCGAA |
| SB8077 | Protospacer (forward) with overhang for ligation into BsmBI-linearized pSB3218 to generate pSB3465 for insertion of *BUB3* upstream of *STU1* using CRISPR. | GACTATTATTGGTCTCATTGTTGA |
| SB8078 | Protospacer (reverse) with overhang for ligation into BsmBI-linearized pSB3218 to generate pSB3465 for insertion of *BUB3* upstream of *STU1* using CRISPR. | AAACTCAACAATGAGACCAATAAT |
| SB8214 | Forward primer to sequence s*tu1-T1080A* CRISPR mutagenesis | ATCCGACGATGCTGTAGACG |
| SB8215 | Reverse primer to sequence *stu1-T1080A* CRISPR mutagenesis | GACCCATATTCTCGTCCCCG |
| SB8243 | Forward primer to mutagenize Stu1 at T1034 to alanine. | GCTATGATTAATCCCTTCAAAAACTTGG |
| SB8244 | Reverse primer to mutagenize Stu1 at T1034 to alanine. | CATTTCCATGCATTTCTTAACATCATTTTCG |
| SB8411 | Forward primer to amplify *SLK19-3xFLAG* | GTGGTAGGAACTTCGTTCATTTTGAGATCCGGGTTTTTCTCC |
| SB8412 | Reverse primer to amplify *SLK19-3xFLAG* | AATTTATGGACATATTGTCGGGTGGAGCTCTAAGCAAATAGC |
| SB8430 | Forward primer to amplify *STU1-3xV5* with homology to pET21b backbone for Gibson assembly. | CTTTAAGAAGGAGATATACAATGTCGTCCTTCAACAATGAGACC |
| SB8431 | Reverse primer to amplify *STU1-3xV5* to add 6xHis tag with homology to pET21b backbone for Gibson assembly. | GGGGTTATGCTAGTTATTGCTCAGTGGTGATGGTGATGATGGCACTGCTCGAGAGCTGTACTATCC |
| SB8438 | To amplify and linearize pET-21b backbone for Gibson assembly | TGTATATCTCCTTCTTAAAGTTAAAC |
| SB8439 | To amplify and linearize pET-21b backbone for Gibson assembly | GCAATAACTAGCATAACCCC |

| | | |
|---|---|---|
| SB8440 | Forward primer to mutagenize Stu1 at T1034 to alanine. | GCTAAAATAGTAAGTGTTTATCAACTGGACC |
| SB8441 | Reverse primer to mutagenize Stu1 at T1034 to alanine. | CATTTCTGAAAGCTTTTTATCTTTGTC |
| SB8450 | To amplify the C-terminus of stu1-2A to generate pSB3564 | TCTAGAGAAAGTTCTGTAAGCTTCACTCC |
| SB8462 | To amplify pSB3519 to remove all *STU1* domains except CL and D4. | TCTAGAGAAAGTTCTGTAAGC |
| SB8463 | To amplify pSB3519 to remove all *STU1* domains except CL and D4. | CATTGTATATCTCCTTCTTAAAGTTAAAC |
| SB8536 | To remove 6xHis tag from Stu1 C-terminus in pSB3519. | GCACTGCTCGAGAGCTGTACTATCC |
| SB8537 | To remove 6xHis tag from Stu1 C-terminus in pSB3519. | TGAGCAATAACTAGCATAACCCC |
| SB8538 | To generate 6xHis tag at Stu1 N-terminus. | ATGCATCATCACCATCACCACTCTAGAGAAAGTTCTGTAAGC |
| SB8544 | Forward primer to sequence *stu1-T1034A* and *stu1-2A* CRISPR mutagenesis | AACTTGAAAAGAAGGATGCCAACTTTAAGG |
| SB8584 | Forward to amplify pSB3554 (pFA6a-link-ymYPET:URA3) to insert 2xymYPET. | ATGGTCTCCAAGGGCGAAGAATTG |
| SB8585 | Reverse to amplify pSB3554 (pFA6a-link-ymYPET:URA3) to insert 2xymYPET. | TTAATTAAACCAGCACCGTCACCGATC |

| | | |
|---|---|---|
| SB8586 | gBlock containing synonymous codon-scrambled 2x mYPet genes separated by 2xGSS linkers, an upstream MCS, and 20 bp homology for Gibson assembly | GACGGTGCTGGTTTAATTAAGGTAC CGGGCCCTCGGTGTAGGATCCTCT AGAATGGTGAGTAAAGGTGAGGAGC TATTTACAGGAGTCGTGCCTATATTA GTTGAACTGGATGGAGACGTTAATG GTCATAAATTTTCAGTATCGGGAGAA GGCGAGGGTGACGCAACATACGG CAAATTAACGCTCAAACTCCTGTGTA CAACTGGTAAGCTACCGGTTCCTTG GCCCACTCTTGTGACTACCTTAGGA TACGGTTTGCAATGCTTCGCAAGGT ATCCCGACCACATGAAGCAACACG ACTTTTTCAAGAGTGCAATGCCTGAA GGATATGTCCAGGAGAGGACAATTT TCTTTAAAGACGATGGTAACTATAAG ACCCGGGCTGAGGTAAAGTTCGAA GGCGATACGCTAGTAAACCGTATTG AGCTAAAGGGAATCGATTTCAAGGA AGATGGAAACATTTTGGGTCACAAAT TGGAATATAACTATAATAGCCATAAT GTGTACATCACAGCCGATAAGCAGA AAAATGGTATAAAAGCTAACTTCAAG ATTCGTCATAACATTGAGGACGGTG GAGTTCAATTGGCCGACCATTACCA GCAAAACACCCCAATTGGTGATGGT CCTGTGTTGTTACCAGATAACCACTA TCTCTCATACCAGTCAAAGTTATCAA AAGATCCTAATGAAAAGCGAGATCA CATGGTCCTGCTAGAATTCCTAACA GCCGCTGGCATTACGCTGGGGATG GACGAGTTGTACAAAGGTTCTTCAG GATCTTCAATGGTTTCTAAAGGTGAA GAGCTATTCACCGGCGTTGTTCCAA TTTTGGTTGAATTGGACGGTGATGTC AACGGGCATAAATTCAGCGTCTCAG GTGAGGGGGAAGGAGACGCTACCT ACGGAAAGTTAACTTTGAAACTTCTC TGTACAACAGGAAAACTACCTGTGC CATGGCCTACGTTAGTCACAACACT AGGTTATGGCTTGCAATGTTTTGCAA GATATCCAGACCATATGAAGCAACA CGATTTTTTTAAGTCCGCTATGCCTG AAGGTTACGTCCAAGAGAGAACCAT TTTCTTTAAAGATGATGGAAATTATAA |

| | | |
|---|---|---|
| | | AACGAGAGCCGAAGTCAAGTTTGAAGGTGACACGCTCGTCAATAGAATAGAGTTGAAGGGCATTGATTTTAAAGAAGATGGTAACATCTTAGGTCACAAGCTAGAGTACAACTACAATTCGCACAATGTCTACATTACTGCTGATAAACAGAAGAACGGTATCAAGGCGAATTTCAAAATTCGACATAACATCGAAGATGGTGGCGTGCAATTGGCCGATCACTATCAACAGAATACTCCTATTGGTGACGGACCCGTATTATTACCCGATAATCATTACCTATCCTATCAATCCAAATTGTCAAAAGATCCAAATGAGAAGCGTGATCACATGGTTTTACTAGAGTTCTTGACTGCTGCCGGCATTACCCTCGGAATGGATGAGCTTTACAAGGGATCTAGTGGTAGTTCTATGGTCTCCAAGGGCGAAGA |
| SB8599 | Forward to amplify *MTW1* with homology to BamHI-linearized pSB3561 for Gibson assembly | GGGCCCTCGGTGTAGATGTCTGCTCCCACTATGAGATCC |
| SB8600 | Reverse to amplify *MTW1* with homology to BamHI-linearized pSB3561 and a linker extension for Gibson assembly | CTCACCATTCTAGAGGATCCTGATGATCTTAACACATCATCAAGTAAATCCAA |
| SB8601 | To amplify pSB3560 backbone to generate *stu1-C-2A*. | AGAACTTTCTCTAGAGTGGTGATGG |
| SB8602 | To amplify pSB3560 backbone to generate *stu1-C-2A*. | ATCGCCTCAAACGAACGGATCC |
| SB8608 | Reverse primer to sequence *stu1-T1034A* and *stu1-2A* CRISPR mutagenesis | CTTTTGACCCATATTCTCGTCCCC |
| SB8714 | gBlock for CRISPR homology-directed repair to generate *stu1-2A (T1034A + T1080A)* | AAGGTAGAAGACATCATTTCTAGAGAAAGTTCTGTAAGCTTCACTCCCATCGACAATAAAAAATCTGAAGGGGATGAGGAATCCGACGATGCTGTAGACGAAAATGATGTTAAGAAATGCATGGAAATGGCTATGATTAATCCCTTCAAAAACTTGGAAACTGATAAAACACTAGAGTTGAAGAATAACGTTGGAAAAAGAACATCAAGCACAGACAGCGTAGTTATT |

| | | |
|---|---|---|
| | | CATGATGATAATGACAAAGATAAAAA GCTTTCAGAAATGGCTAAAATAGTAA GTGTTTATCAACTGGATCAGCCAAA CCCAGCAAAGGAGGAAGATGATATA GATATGGAAAATTCTCAAAAATCTGA TTTGAATTTAAGTGAAATTTTTCAAAA CAGTGGTGAAAATACCGAAAGGAAA TTGAAGGACGATA |
| SB8715 | gBlock for CRISPR homology-directed repair to generate *stu1-T1034A* | AAGGTAGAAGACATCATTTCTAGAGA AAGTTCTGTAAGCTTCACTCCCATC GACAATAAAAAATCTGAAGGGGATG AGGAATCCGACGATGCTGTAGACG AAAATGATGTTAAGAAATGCATGGAA ATGGCTATGATTAATCCCTTCAAAAA CTTGGAAACTGATAAAACACTAGAGT TGAAGAATAACGTTGGAAAAAGAAC ATCAAGCACAGACAGCGTAGTTATT CATGATGATAATGACAAAGATAAAAA GCTTTCAGAAATGACGAAAATAGTAA GTGTTTATCAACTGGATCAGCCAAA CCCAGCAAAGGAGGAAGATGATATA GATATGGAAAATTCTCAAAAATCTGA TTTGAATTTAAGTGAAATTTTTCAAAA CAGTGGTGAAAATACCGAAAGGAAA TTGAAGGACGATA |
| SB8716 | Forward to amplify gBlocks SB8714 and SB8715 for CRISPR transformation. | AAGGTAGAAGACATCATTTCTAGAG |
| SB8717 | Reverse to amplify gBlocks SB8714 and SB8715 for CRISPR transformation. | TATCGTCCTTCAATTTCCTTTC |
| SB8727 | To amplify *MPS1* to insert into pCDFDuet1 containing Lambda PPase (pSB3587) | CATCACCACAGCCAGATGTCAACAA ACTCATTCCATG |
| SB8728 | To amplify *MPS1* to insert into pCDFDuet1 containing Lambda PPase (pSB3587) | CCGAGCTCGAATTCGCTAAATTTTGT AATCTGCAAATTTCC |
| SB8762 | Forward to amplify gBlock SB8764 for CRISPR mutagenesis. | TACCATGTTATTTTAATGCGTTACC |

| | | |
|---|---|---|
| SB8763 | Reverse to amplify gBlock SB8764 for CRISPR mutagenesis. | GGTACGAACGGTAAGCGTAATG |
| SB8764 | gBlock for CRISPR homology-directed repair insertion of the *BUB3* open-reading frame upstream of *STU1* | TACCATGTTATTTTAATGCGTTACCC<br>CGTCGGCGCGTATTGGCGCATTAA<br>CATCTTTATAGAAAATCAAATTATTTA<br>CTTATTTATTCTAAAACGCTAAACATC<br>TATTTATTTACATTTGAGGTTGAAACTT<br>CCCTTCATCCGATTGACAGGCATAT<br>TTAGCGGTAATTATTAGGGTTTTTGGA<br>GAGACCTTGTATTCTTCAGAAATAAT<br>GCAGATAGTACAAATTGAGCAGGCC<br>CCAAAAGACTACATAAGCGACATCA<br>AAATAATCCCTTCCAAGTCACTGCTT<br>TTGATTACGTCTTGGGATGGCTCTTT<br>AACAGTCTACAAATTCGACATTCAAG<br>CAAAGAATGTTGACCTTTTACAATCG<br>CTACGATATAAACATCCGTTATTGTG<br>CTGCAATTTCATCGACAATACCGAT<br>CTGCAAATATACGTGGGAACTGTAC<br>AGGGTGAAATTCTAAAAGTTGATTTG<br>ATAGGTAGTCCCAGCTTCCAAGCTT<br>TGACGAACAATGAAGCCAATTTGGG<br>TATTTGCCGAATATGCAAATATGGAG<br>ACGATAAACTCATTGCCGCGTCATG<br>GGATGGCCTGATAGAGGTTATCGAC<br>CCTCGCAATTATGGTGATGGAGTTAT<br>TGCTGTTAAAAATTTGAACTCTAATAA<br>CACAAAGGTGAAGAATAAGATATTTA<br>CTATGGATACAAACTCCTCTCGATTG<br>ATCGTTGGTATGAACAATAGTCAGGT<br>TCAATGGTTTCGCCTGCCACTCTGT<br>GAGGATGATAACGGAACAATTGAAG<br>AATCAGGACTGAAGTACCAAATAAG<br>AGATGTCGCTCTTTTACCGAAAGAA<br>CAAGAAGGTTATGCATGTAGCAGCA<br>TTGACGGGCGAGTTGCTGTGGAGTT<br>TTTCGATGATCAGGGCGATGATTACA<br>ACTCAAGCAAAAGATTTGCATTTAGA<br>TGCCACCGTTTGAATTTAAAAGATAC<br>AAACTTAGCGTATCCAGTAAATTCTA<br>TTGAATTTTCCCCCCGTCATAAGTTC<br>CTATACACGGCTGGCTCTGATGGCA<br>TAATTTCATGCTGGAACCTACAAACC |

| | | |
|---|---|---|
| | | CGCAAGAAAATAAAAAATTTCGCCAAATTTAACGAAGACAGCGTGGTTAAAATTGCTTGTTCGGACAATATTCTATGTCTGGCAACTTCTGATGATACTTTCAAGACAAACGCCGCAATTGACCAAACTATTGAACTAAACGCAAGTTCAATATACATAATATTTGACTATGAGAACGGTTCTTCAGGATCTTCCGGTTCCAGTATGTCGTCTTTCAACAATGAGACCAATAATAACAGCAACACTAATACACATCCAGACGATTCCTTCCCTTTGTATACCGTATTCAAGGACGAGTCTGTACCCATCGAGGAAAAAATGGCACTGCTTACACGGTTCAAAGGACATGTAAAAAAGGAACTAGTTAACGAATCGTCGATCCAAGCTTATTTCACTGCGTTACTGTTCATCTCCGGCCATTACGCTTACCGTTCGTACC |
| SB8767 | To remove N-terminus of Mps1 in pSB3593 | AATAATAGAAATATAATTACAGTAAATGACTCC |
| SB8768 | To remove N-terminus of Mps1 in pSB3593 | CTGGCTGTGGTGATGATG |
